# The role of cognition and movement of seed-dispersing animals in shaping plant distributions

**DOI:** 10.1101/2023.08.01.551244

**Authors:** Benjamin Robira

**Affiliations:** Animal Ecology Unit, Research and Innovation Centre, Fondazione Edmund Mach, San Michele all’Adige, TN, Italy

**Keywords:** Agent-based model, Feedback loop, Foraging, Frugivory, Memory, Movement, Phenology, Zoochory

## Abstract

In the scenarios concerning the emergence and selection of spatiotemporal cognitive abilities in vagile plant-eating animals, there is always an implicit assumption: the distribution of plants does not change and ultimately shapes the cognitive abilities of the animals, hence their movement. Yet, if plant distribution patterns are likely to remain unchanged over short time periods, they may change over long time periods as a result of animal exploitation. In particular, animal movement can shape the environment by dispersing plant seeds. Using an agent-based model simulating the foraging behaviour of a seed disperser endowed with spatiotemporal knowledge of resource distribution, I investigated whether resource spatiotemporal patterns could be influenced by the level of cognition involved in foraging. This level of cognition represented how well resource location and phenology were predicted by the agent. I showed that seed dispersers could shape the long-term distribution of resources by materialising the routes repeatedly used by the agent with the newly recruited plants. This stemmed from the conjunction of two forces: competition for space between plants and a seed-dispersing agent moving from plant to plant based on spatiotemporal memory. In turn, resource landscape modifications affected the benefits of spatiotemporal memory. This could create eco-evolutionary feedback loops between animal spatiotemporal cognition and the distribution patterns of plant resources. Altogether, the results emphasise that foraging cognition is a cause and a consequence of resource heterogeneity.

## I Introduction

**H**ypotheses about the evolution of vagile plant-eating foragers’ spatiotemporal cognition (i.e., the ability to process, integrate, and utilise information on resource distribution and phenology) generally assume that foragers evolve in an unchanging environment (Rosati, 2017). However, while foragers’ cognition is likely to be shaped by the distribution of resources (Boyer & Walsh, 2010; Bracis et al., 2015; Grove, 2013; Milton, 1981; Robira et al., 2021), the characteristics of the resources themselves could be the result of forager-induced selection (Lomáscolo & Schaefer, 2010; Rojas et al., 2021; Soldati et al., 2015). In the tropical rainforests, for example, the cognition of frugivorous primates has probably been shaped by the challenge of the ephemerality and dispersion of their food (Milton, 1981; Trapanese et al., 2019a; Zuberbühler & Janmaat, 2010), supporting complex (Janmaat et al., 2013; Janson, 1998, 2016; Robira et al., 2023a; Teichroeb & Vining, 2019; Trapanese et al., 2019b) and versatile (Janmaat et al., 2006; Robira et al., 2023b; Trapanese et al., 2022) movement heuristics. In turn, primates remember and forage for the most rewarding plants (Ban et al., 2014; Ban et al., 2016; Flörchinger et al., 2010). The most attractive plants are thus more likely to have their seeds dispersed, driving plant and landscape evolution [soldati2017long; Chapman et al. (2013); Lambert and Garber (1998)].

The way seeds are dispersed contributes to the emerging pattern of resource distribution (Vittoz & Engler, 2007). Most of the world’s trees (Tiffney & Mazer, 1995) and especially rainforest trees (Bagchi et al., 2011; Gottsberger & Silberbauer-Gottsberger, 1983) are dispersed by animals (zoochory). Compared to other modes of dispersal, such as wind (anemochory) or gravity (barochory), zoochory induces a less patchy distribution of adult trees (Seidler & Plotkin, 2006; but see Hubbell, 1979). However, animal-dispersed fruit trees are still not homogeneously distributed (Hubbell, 1979). This is because although travelling long distances, animals may tend to revisit memorised sites regularly, if not exclusively (Addis et al., 2017; John et al., 2016; Soldati et al., 2015). Furthermore, the imperfection and limited size of spatial memory (Fagan et al., 2013) is expected to lead to different movement patterns (Avgar et al., 2013), and therefore potentially to different patterns of resource distribution (Côrtes & Uriarte, 2013).

By influencing the spatial distribution of plants, animals may also shape the relationship between the spatial and temporal availability of resources. Indeed, the tendency of animals to repeat travel decisions (i.e., their routine, Riotte-Lambert et al., 2017), may result in seeds being deposited along fixed spatial sequences. Thus, all trees fruiting at the same time may be deposited in the same place, affecting their local synchrony in food production. Synchrony in productivity can then be used by animals to optimise movement decisions (Janmaat et al., 2012). By being affected by cognition and by affecting resource dispersal, animal movement could thus lead to eco-evolutionary feedback loops between animal cognition and resource distribution (Borah & Beckman, 2022). In other words, foraging cognition could be both a cause and a consequence of resource heterogeneity.

To illustrate the possible retroactions between cognition and resource distribution, let’s consider an extreme speculative scenario in which seed-dispersing frugivores are freely foraging (Figure 1). At start, assuming a random homogeneous distribution, long-term spatiotemporal memory can be advantageous and selected for (Bracis et al., 2015; Grove, 2013). Through zoochory, deliberate targeting of some resource patches over others may induce a change in resource distribution over time (e.g., increasing patchiness and local synchrony among trees), eventually making cognition more and more advantageous (Boyer & Walsh, 2010; Bracis et al., 2015; Grove, 2013). Gradually, by repeatedly targeting the same plants, the routes used by the foragers may begin to materialise through the seeds scattered along the way (Di Fiore & Suarez, 2007). However, in doing so, foragers who can only passively follow these routes due to their sensory abilities and have survived may also begin to become very efficient: they should not even have to bear the cost of high cognitive ability, putting them at an advantage over foragers who bear the burden of cognition (Burns et al., 2011; Raichle, 2006). While the pattern of resource distribution can be maintained because the routes are used equally by all foragers (but through two different mechanisms: long-term memory *vs* taxis/kinesis, Benhamou & Bovet, 1992; Benhamou & Poucet, 1995), cognition could even start to be counter-selected.

**Figure 1.**
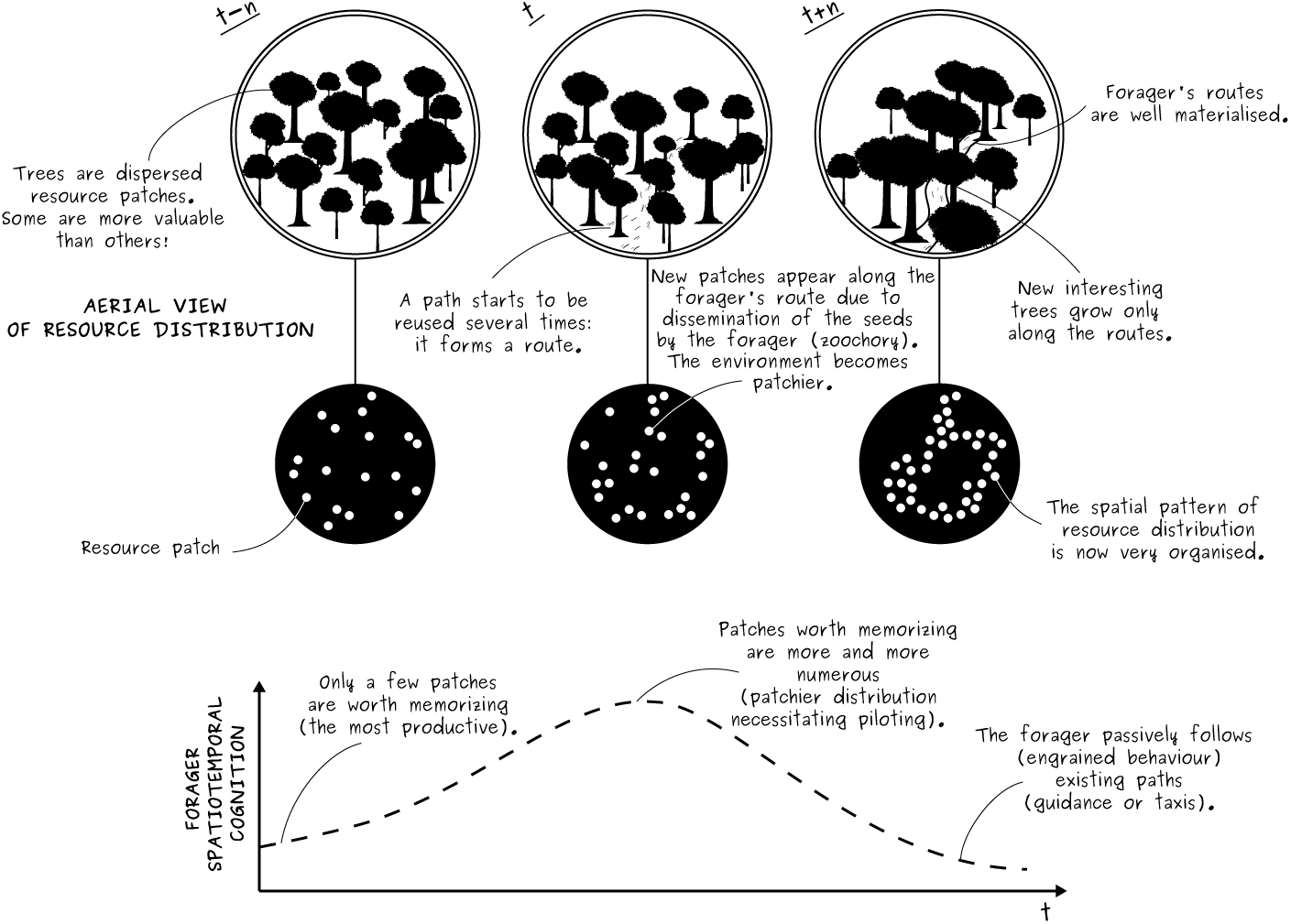
Theoretical expectations on eco-evolutionary feedback loops between animal spatiotemporal cognition and resource distribution

While some facets of this scenario may be rooted in empirical evidence (e.g., forest elephant paths matching with food distribution, Blake et al., 2009; Blake & Inkamba-Nkulu, 2004; Campos-Arceiz & Blake, 2011), most of it remains speculative. The aim of this study is to put such a scenario to the test using an agent-based model simulating the movement of seed-dispersing frugivores (Figure 2). Specifically, I aim to identify the conditions that favour the emergence of an eco-evolutionary feedback loop between animal spatiotemporal cognition and resource distribution. To do this, I first varied two forager traits (the amount of knowledge about plant location and phenology, and movement rules) and one plant trait (spatial competition) to measure how this affected plant aggregation (patchiness), the formation of materialised routes by the plants (alignment), and the local synchrony of plant phenology (spatial autocorrelation in fruiting dates) after the forager foraged for a long time and seed dispersal occurred. I predicted that higher spatiotemporal knowledge would lead to the emergence of route-like patterns (see Appendix, Using patchiness and alignment to characterise spatial point patterns for how this is evidenced by resource patchiness and alignment values) with locally synchronous fruiting as a consequence of routine movements. As animals may show different levels of opportunistic exploration *en route* to the target (e.g., stopping by to all plants, or heading directly to the target), I tested whether the magnitude of changes varied with the agent movement rules (i.e., if the agent stopped at all plants encountered on the way to the target plant, rather than passing all or only non-fruiting other perceived plants). In addition, as plants may also play a role in their own distribution (e.g., through competition for space, Craine & Dybzinski, 2013), I repeated these scenarios considering different level of spatial competition between plants to test whether spatial competition could buffer (expected for high competition) or promote (expected for low competition) the observed changes induced by the forager’s different cognitive levels. Second, I investigated whether variations in space engineering would affect the forager’s foraging efficiency (i.e., the ability to find a lot of food by travelling a short distance). Following the illustrative aforementioned scenario, I expected that engineering should favour cognition (i.e., higher foraging efficiency) up to an extreme case (the formation of materialised travelling routes).

**Figure 2.**
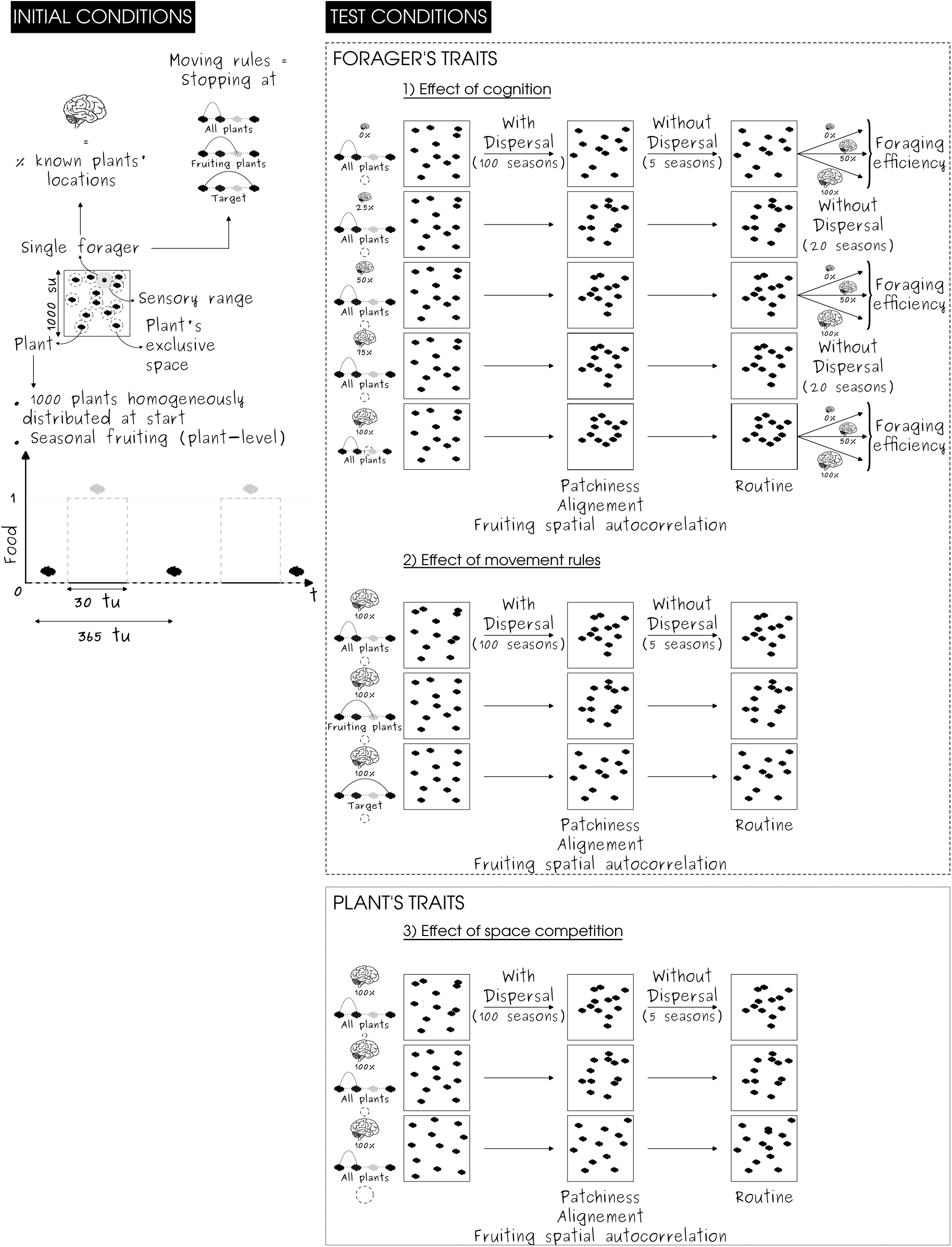
Illustration of the model workflow

## II Material and Methods

### II.1 Agent-based model

The main objective of the model was to simulate the foraging behaviour of a single agent in an environment that is dynamical both on the short term, through an interplay between resource growth and depletion, and on the long term, due to seed dispersal (Figure 2). For simplicity, I will refer to the agent as the forager, and to resource patches as fruit plants. The arbitrary units for both space and time makes this model a simplified representation of possibly a wide variety of systems, depending on the set of parameters chosen. For example, assigning a spatial unit of one metre, a temporal unit of one day, and low spatial competition between plants could be consistent with an asynchronous tropical tree forest in which a large terrestrial mammal forages. The model was implemented in the *C++* language in *R* software (v.4.2.1, R Core Team, 2022) based on the *Rcpp* package (Eddelbuettel, 2013; Eddelbuettel et al., 2023; Eddelbuettel & François, 2011). The model parameterisation is available in Table 1.

**Table 1:** Model parameters.

| Modelling entity | Parameter | Definition | Value | Unit |
| --- | --- | --- | --- | --- |
| Environment | Map size | Length of a side of the square environmental map | 1000 | su |
| Environment | Quadrat size | Length of a side of a square quadrat to calculate Lloyd index of patchiness | 50 | su |
| Environment | Period length | Length of a period before a given plant starts producing again | 365 | tu |
| Environment | Number of seasons | Number of seasons (with seed dispersal plus without seed dispersal) before the simulation is ended | 100 + 5 | - |
| Environment | Number of plants | Number of plants hosted by the environment | 1000 | - |
| Environment | Fruiting length | Time duration of the fruiting period of each plant | 30 | tu |
| Environment | Maximum food yielded at a plant | Food quantity that a plant might yield at best (peak of the triangular-shaped food distribution) | 1 | fu |
| Environment | Spacing intensity | Relative length of a square map whose area would correspond to the area of exclusive spaces of all plants without overlapping | 5, 45, 85 | - |
| Agent | Speed | Speed at which the forager moves | (500,1000,2000) | su/tu |
| Agent | Torpor time | Time duration for which the forager stops foraging in case no food is available in the environment | 1 | tu |
| Agent | Perceptual range | Distance at which the forager is aware of the environment | 15.81 | su |
| Agent | Knowledge rate | Proportion of plants of the environment for which the forager knows the location and production timing | (0, 0.25, 0.5, 0.75, 1) | - |
| Agent | No-return time | Time delay before a forager mentally decides to target a previously visited plant | 2 | tu |
| Agent | Dispersal time | Time duration during which seeds from a previously ingested fruit can be dispersed | 0.5 | tu |
| Agent | Probability of dispersal | Probability (per tu) that the seeds is actually dispersed | 0.02 | 1/tu |
| Agent | lambda["step length"] | Average step length for random movements used to parameterise the exponential distribution | 0.01 | su |

#### Environment

##### Spatial distribution

I modelled the environment as a flat square map of side 1000 arbitrary spatial units (su). Within this map, 1000 plants were initially homogeneously distributed (uniform distribution of their *x* and *y* coordinates, resulting in a Poisson distribution of plant density).

##### Resource growth

Time was cyclical, with a season of 365 arbitrary time units (tu). Each plant produced resource (fruit containing seeds) for 30 tu only once every season. Periods of productivity of each plant were randomly distributed (i.e., uniform distribution of the start date). Thus, resource was seasonal at the plant-level, but not at the map-level. Productivity was characterised by a triangular-shaped pattern of the quantity of food over time: at start of the productive period, food quantity started to increase linearly up to 1 arbitrary food unit (fu), mimicking progressive fruit ripening, and then decreased linearly to 0 fu, mimicking fruit rotting. The depletion of the food by the forager had no effect on this dynamic. Food could continue to ripen (though never to the same extent as if it had not been depleted) and rot. At a given time, the quantity of food available at a given plant thus depended on the productivity of the plant at that time, as well as the depletion history by the forager, creating a short-term dynamics of resource temporal distribution.

##### Seed dispersal

Trees reproduced by ingestion of ripe fruit by the forager and subsequent dispersal. Once the forager ate ripe fruit, seeds could be dispersed only one time up to 0.5 tu after ingestion. Dispersal occurred at a probability of 0.02 tu^*−*1^, independent of the quantity of fruit ingested. This probability was considered low, to mimic the natural slow pace of land use changes. All fruit eaten did not mandatorily lead to dispersal. Dispersal was eventually successful only if some space was left available for the new plant: no plant had to be at a distance less than 3.61 su. This number was obtained such that the cumulative (when non-overlapping) exclusive “lifespace” by plants (a circular area of radius the aforementioned distance) could reach the area of a map of length 0.45 times that of the full map (but other values, referred as spacing intensity were tested, see Route formation as a product of two forces). If it happened that there was available space for the seedling to grow, then the seedling location was sampled in the available space along the linear path linking the previous and current locations. Otherwise, the seedling could not grow and died.

If the seedling was successfully dispersed and recruited, an old plant (already present in the map) was randomly selected and died to keep density constant. The start date of fruiting of the newly recruited plant was randomly sampled within a reduced Gaussian distribution (i.e., variance of one) centered on the start date of fruiting of the parent plant. Because memory size can affect foraging efficiency, to keep it constant, the location and fruiting time of the newly recruited plant were memorised if it replaced a plant that was part of the forager’s memory (see Appendix, Sensitivity to the learning pathways following old plant mortality for a test of an alternative learning way). The newly recruited plant was considered as mature only the season after. Seed dispersal could thus induce a long-term dynamics of resource spatial distribution and of the distribution of starting dates in the plant population.

#### Forager

##### Cognitive abilities

The forager was endowed with a gradient of knowledge abilities: from no memory of the locations of plants (naive forager) and the timing of their period of productivity to perfect omniscience (i.e., all plant locations and associate phenology are known). Otherwise, it only knew a certain proportion (fixed for the whole simulation) of plant locations and their associated phenology (hereafter *spatiotemporal knowledge rate*). These plants were selected randomly at the start of the simulation. When a plant was part of the forager’s memory, the forager knew both the spatial (location) and temporal (fruiting timing) attributes. Thus, it could not know only one or the other.

##### Movement rules

The forager was initially randomly located within the map. It is fully aware of plants present within sensory range fixed to 15.81 su. This actually corresponds to the average nearest-neighbour distance which equates 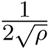, with *ρ* the density of plants (here 0.001) at start of the simulation where plants are homogeneously distributed. It is also aware of plants that are part of its spatiotemporal long-term memory. To decide where to head next, the forager could either rely on its private knowledge (sensory or memory) or not. If all plants within its private knowledge did not yield any fruit (i.e., all having less than 0.001 fu), the forager was moving randomly. Random movements were performed by randomly choosing a direction (uniform distribution of the heading angle), while the step length was sampled within an exponential distribution of mean *λ_step length_* = 0.01. An exponential distribution of step lengths is indeed classically observed in nature (Benhamou, 2014). Otherwise, the forager attempted to maximise its predicted short-term foraging efficiency (i.e., of the next movement step) and targeted the closest plant which maximised it. Predicted short-term foraging efficiency was defined as the ratio between the move benefits, here the predicted food quantity at the plant (which accounts for depletion history), and the move costs, here linearily proportional to the distance to the plant. Nonetheless, the forager avoided to target plants recently visited within the previous 2 tu to avoid targeting recently depleted plants (Robira et al., 2021).

Once the target had been chosen (or random point, if none), the forager moved straight to the first plant that should have been theoretically perceived *en route* to the initial destination point if any (provided they were not recently visited), or the destination otherwise. When this destination was reached, it depleted all the food available and then re-estimated the best plant to target and so on. After each moving bout time was updated based on the distance travelled, knowing that the forager moved at a speed of 1000 su/tu (see Appendix, Sensitivity to the agent’s speed for results with an agent moving at half, or twice, this speed). Then, the resource quantity available at each plant was updated.

In case all plants were empty (no food available in the environment, considered if the sum of food available was less than 1 fu), the forager entered a torpor period for 1 tu, and so on until food was available again. This was done to avoid biasing measure of foraging efficiency in case the forager could deplete entirely the environment while being unable to disperse to another area, as expected in nature if this happens.

### II.2 Analyses

#### Characterisation of plant distribution patterns

To assess whether different levels of cognition resulted in different organisations of the spatial distribution of plants at the end of the simulation, I focused on three metrics describing plant spatiotemporal patterns: the *patchiness*, the *alignment* and the *spatial autocorrelation of fruiting timing*.

##### Patchiness

Patchiness (i.e., the tendency of plants to aggregate into clusters) was measured using Lloyd’s index of patchiness (Lloyd, 1967). This index is given by the following formula

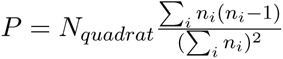

where *n_i_* represents the number of plants in quadrat *i* and *N_quadrat_* the number of quadrats. The map was divided in 400 square quadrats of side 50 su. The quadrat size, provided sufficient length for biological relevance, little affects the measurement (Lloyd, 1967). Lower values indicate homogeneous distribution while larger values indicate heterogeneous and route distributions (see Appendix, Using patchiness and alignment to characterise spatial point patterns).

As the forager is set up to forage in a square map with reflective boundaries, seed dispersal may lead to the shrinkage of resource range towards the centre of the map (where recursions are inevitably higher). As patchiness is estimated on the whole map, this shrinkage would contribute to inflating the calculated patchiness. Indeed, patchiness measures the tendency of plants to have more neighbours in their vicinity (i.e., quadrat size) than expected based on the density in their overall range. By affecting this range, shrinkage may lead to more ‘empty’ patches and more ‘crowded’ plant patches, but only because the area covered by plants has been reduced (forming one large patch, an artefact not of interest here as the forager will limit foraging in this large patch), not because plants have been regrouped into multiple patches (which is of interest here as this affects the benefits of cognition). I therefore calculated a measure of shrinkage *s* as one minus the area covered by the 95% of the Gaussian-based kernel distribution based on all plant locations divided by the map size (“kernelUD” function of the *adehabitatHR* package (Calenge, 2006), with a smoothing parameter *h* set to 50; the kernel was bounded by the map borders). To account for variations in shrinkage in the different simulated scenarios (see Appendix, Variations in resource range shrinkage intensity), I corrected patchiness such as *P_c_* = *P* (1 *− s*). In this way, for a similar spatial distribution, patchiness will be more important if the overall range is large, that is, if the shrinkage is low.

##### Plant alignment

Routes are (curvi-)linear features. To assess the tendency of plants to form linear motifs, I quantified their alignment. For a given plant, alignment *a* was defined as one minus the minimum of the sine of the angles of the triangle formed by the plant of interest and the two closest plants. This varies from perfect alignment (value of 1), to no alignment (i.e., the minimum angle would be at worst 60° hence an alignment of ca. 0.134). Therefore, I used a corrected alignment *a_c_* varying from 0 (no alignment) to 1 (perfect alignment) such as

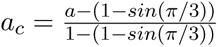

An environment where routes are formed should result in many points having a high alignment (value close to one) and fewer low values (close to zero, such as the hubs of the route network); this means that high values should be over-represented compared to low values, resulting in a long left tail, also known as negative skew. To have an index ranging from 0 (low alignment) to 1 (high alignment), I defined the general alignment for a given environmental map as the opposite of the skewness of alignment values of all plants in the map. In this case, positive values indicate the existence of linear motifs while negative values indicate no obvious linear motifs. As confirmed from simulations, random distributions (homogeneous or heterogeneous) should show moderate positive values, while a route pattern should have a high positive value (see Appendix).

##### Spatial autocorrelation of fruiting timing

Spatial autocorrelation of fruiting timing (i.e., the tendency of fruiting date of plants to more closely match that of neighbours than of distant plants) was calculated as the Moran Index, *I*, applied to the start of fruiting dates. Since the “start of fruiting dates” corresponds to a circular variable, I followed (Schmal et al., 2017) to adapt existing functions of the *ape* package (Paradis & Schliep, 2019) to circular variables. Moran’s *I* was therefore defined as

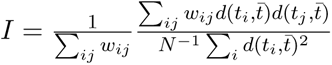

where *N* is the number of plants, *t* is the fruiting date (transformed to radian), and 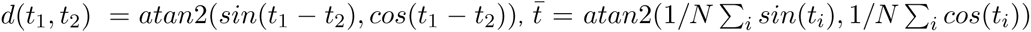, that is the angular difference between the two circular variables *t*_1_ and *t*_2_. I used the inverse of the Euclidean distance between plant locations as weights *w*. Values of *I* close to 0 indicate absence of spatial autocorrelation. Values close to -1 indicate negative spatial autocorrelation. Values close to 1 indicate positive spatial autocorrelation.

##### Routine emergence

For each simulation, I verified whether a routine behaviour emerged and paralleled the variations in plant distribution. Routine was defined as the predictability of the plant visit sequences during five supplementary seasons in which dispersal could not occur, such as the spatial distribution (and plant id) remained unchanged. Predictability was defined as one minus the first-order conditional entropy (*H*_1_) of the plant id visit sequence (Riotte-Lambert et al., 2017). *H*_1_ characterises the average uncertainty about the next plant to be visited (*T_v_*), knowing the previous visit *T_p_* (Riotte-Lambert et al., 2017), such as: 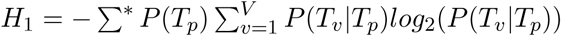 where 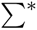 represents all possible *T_p_*, *V* the number of visits, and *P*() means “probability of”. I computed the routine possibly up to only a first-order conditional entropy, as computing routine indices is computationally demanding and a first order is expected to be the most informative to describe a perfect traplining behaviour (Riotte-Lambert et al., 2017). The routine index varies from 0 (no routine) to 1 (perfect traplining). I computed the routine possibly up to only a first-order conditional entropy, as computing routine indices is computationally demanding and a first order is expected to be the most informative to describe a perfect traplining behaviour (Riotte-Lambert et al., 2017). The routine index varies from 0 (no routine) to 1 (perfect traplining).

#### Research questions and associated scenarios

##### Q1. Does resource spatiotemporal distribution vary with cognition level?

To investigate whether different levels of cognition eventually led to different distribution patterns, I considered five forager types differing by the proportion of plants for which the location and phenology were known (from 0 to 100%, at a step of 25). For each forager, 200 simulations were performed for a given condition (i.e., spatiotemporal knowledge rate, constant throughout the entire simulation), in which the forager foraged individually 100 seasonals.

##### Q2. Does resource distribution engineering favour some cognition levels?

To investigate whether the engineered distribution of plants could in turn affect the foraging benefits purported by cognition, I used the final plant distributions of simulations computed to answer question 1). For each of these plant distributions, in three different simulation runs, I let forage a forager endowed with no, intermediate (i.e., half of plants known), or full (all plants known, omniscience) spatiotemporal knowledge. In these simulations, no seed-dispersal could occur. I measured the foraging efficiency after the forager foraged for 20 full seasons and compared them across environments and forager abilities. Foraging efficiency was computed as the cumulated quantity of eaten food, over the cumulated distance travelled. I verified that at the end of the simulation that foraging efficiency stabilised, which meant that I reached a foraging equilibrium. I considered that foraging efficiency stabilised when the foraging efficiency sampled after each moving bout during the fourth and fifth quintile of simulation times did not differ of more than 5% (reference is the minimum efficiency in the fourth or the fifth quintile of simulation times). I performed 200 simulations for a given condition (i.e., knowledge rate).

##### Q3. Route formation as a product of two supplementary forces? Plant competition for space and forager movement step rule

During the model construction, I noticed that two mechanisms could amplify the phenomenon of route formation, in addition to the use of memory: space competition between plants and sequential targeting of all plants seen *en route* to the target. To better quantify the effect of those two forces, I considered two other sets of 200 simulations each.

To assess the effect of competition for space between plants, I compared the final plant spatiotemporal patterns when an omniscient forager (all plant locations and phenology known) was let foraging in the same setting as in question 1, with the only difference that the spacing intensity could be of 5 (small lifespace, high local plant density), 45 (intermediate lifespace, intermediate local plant density and reference setting) or 85% (large lifespace, low local plant density), so that the radius of this exclusive circular lifespace was always inferior to the perception range (12.89 su vs 15.81 su).

To assess the effect of the movement rule of the forager on plant distribution patterns, I compared the final plant spatiotemporal patterns when an omniscient forager was let foraging in the same setting as in question 1) and while the stringency in skipping plants encountered *en route* varied. It could either stop at all plants perceived on the way to the initial target (mimicking a “monitoring” behaviour, as in all other simulations; small skipping stringency), stop only at plants currently fruiting (i.e., with at least 0.001 fu, mimicking an opportunistic feeding only; intermediate skipping stringency) or move directly to the target (large skipping stringency).

## III Results

### III.1 Resource distribution is affected by the forager’s cognition

Using simulated environments (see Appendix), the expected patchiness for route patterns was 1.514 times and 1.166 times superior to homogeneous and heterogeneous distribution patterns, respectively. Alignment was 2.913 times and 3.147 times higher in route patterns than in homogeneous and heterogeneous distribution patterns, respectively. In homogeneous environments, considering high local synchrony in fruiting dates resulted in spatial autocorrelation 23.454 times higher than considering intermediate local synchrony. In heterogeneous environments it was 28.291 times higher than considering intermediate local synchrony. In both homogeneous and heterogeneous environment, Moran’s Index reached a value close to 0.008 for intermediate local synchrony, and was coherently close to 0 when no local synchrony was modelled (see Appendix, Using Moran’s Index to characterise plant synchrony). These benchmarks, useful to calibrate the magnitude of the changes, are shown in (Figure 3).

**Figure 3.**
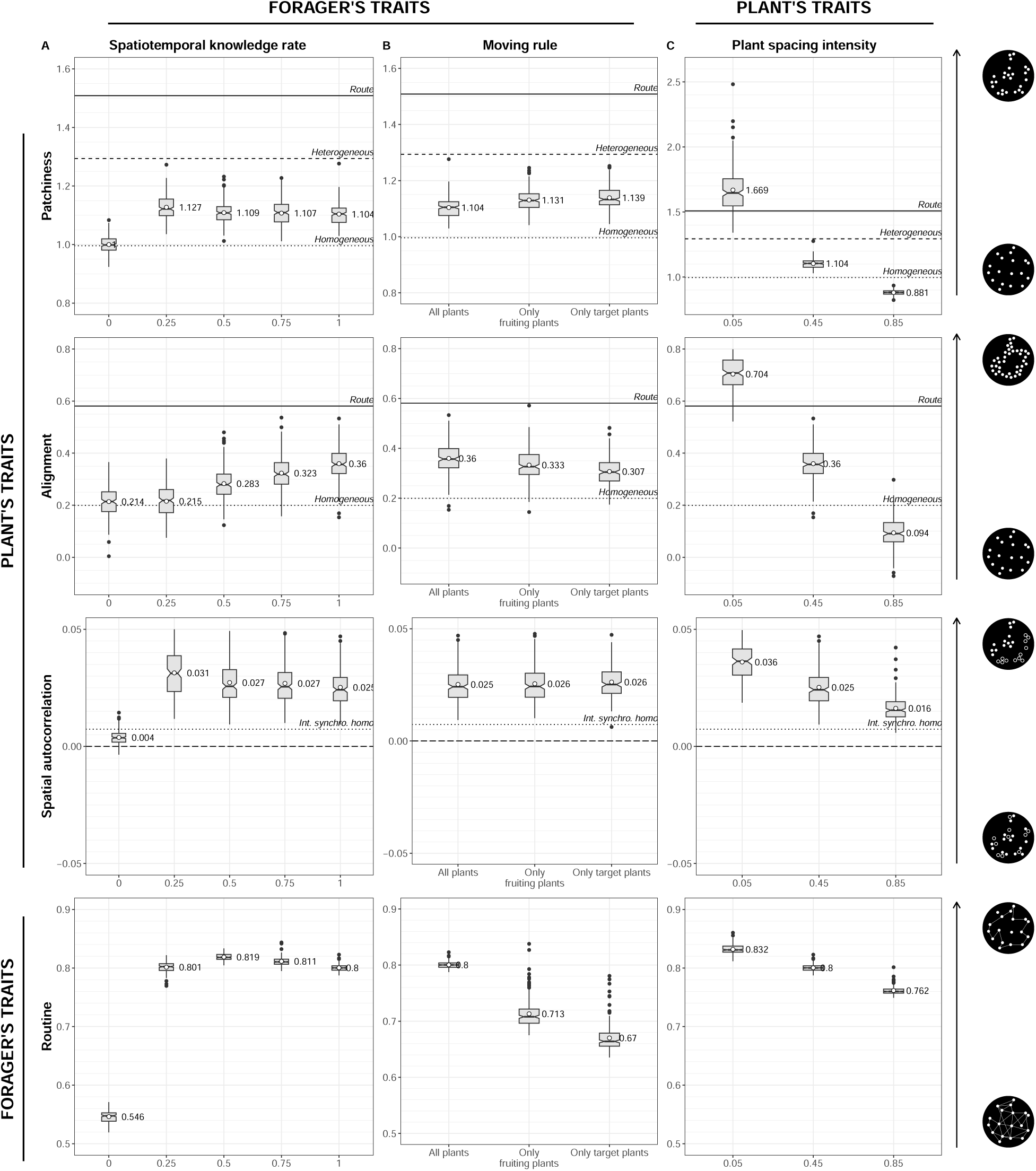
Seed-dispersing foragers can shape resource distribution | The notched box plots show the results of 200 simulations when a seed-dispersing forager with varying level of cognition (i.e., proportion of plant locations and associated phenology known), moving choice rules or space competition intensities between plants (the two latter being simulated with an omniscient forager; left to right) was let foraging for a long time. The resulting distribution pattern was assessed by its patchiness (the tendency of plants to be aggregated into patches; larger values indicate highly aggregated plants), its alignment (the tendency of plants to form linear geometries; larger values indicate the presence of linear geometries) and spatial autocorrelation in fruiting date (the tendency of plants to have fruiting dates closer/farther in time to their spatial neighbours; values close to -1 indicate negative spatial autocorrelation, values close to 0 indicate no spatial autocorrelation, values close to 1 indicate positive spatial autocorrelation). Indices’ benchmarks obtained from simulated environments are represented by the horizontal lines (Homo. = Homogeneous, Hetero. = Heterogeneous, Synchro. = Synchrony) (see Appendix). Because of overlap, only alignment a1n4d intermediate synchrony in a homogeneous (and not heterogeneous) environment was plotted. The emerging movement behaviour of the forager was described in terms of routine behaviour (the tendency of the forager to target plants in a predictible way; values close to 0 indicate no predictible movements, values close to 1 indicate highly predictible movements).

Foragers’ spatiotemporal knowledge rate affected the resource distribution emerging after it had been let foraging for a long time (Figure 3A). Both the patchiness (the tendency of plants to be aggregated into patches) and the alignment (the tendency of plants to form linear geometries) abruptly changed when the forager was cognitively endowed. Specifically, patchiness followed a ratchet effect, abruptly increasing as soon as the forager was cognitively endowed (spatiotemporal knowledge rate *≥* 0.25) up to being 1.199 times superior to when a naive forager foraged. Alignment progressively increased with the forager’s spatiotemporal knowledge, up to being 1.681 times superior to when a naive forager foraged. (Figure 3A). In other words route-like patterns started to emerge as a consequence of memory-driven movements.

Spatial autocorrelation in fruiting date (the tendency of plants to have fruiting dates closer/farther in time to their spatial neighbours) somehow followed the same trend as patchiness (Figure 3A). Specifically, when a forager endowed with spatiotemporal knowledge foraged, the spatial autocorrelation in fruiting dates was about 7.713 times (mean value = 0.029) more important than when a naive forager foraged (with a spatial autocorrelation close to 0). In other words, the synchrony in fruiting among neighbouring plants started to emerge as a consequence of memory-driven movements.

Overall, space engineering was paralleled with the emergence of routine behaviours, which increased with spatiotemporal knowledge rate to saturate around 0.8 (Figure 3A). This held when no dispersal occurred (Figure A6).

These results were robust to the speed at which the agent moved, but were somewhat hampered by the inability of the agents to learn the new seedling locations when an old tree died (instead the agent learned the location and phenology of the last fruiting plant visited; Figure A8).

### III.2 Engineered patterns affects the benefits of spatiotemporal cognition

On average, an “omniscient” forager (knowing all plants’ locations and phenology patterns) was around 1.316 times more efficient than an “intermediate” forager (knowing only 50% of them) or 3.666 times than a “naive” forager (knowing none of them) (Figure 4). While the benefits of omniscience relatively to a naive forager were rather constant (min-max = 3.582-3.737 times larger), the relative benefits of intermediate memory varied a little with the resource distribution emerging from previous engineering (Figure 4). Compared to when the foragers foraged in an environment previously shaped by a naive forager (left boxplots in Figure 4), the relative foraging efficiency of intermediate foragers compared to naive foragers increased of 5.7% and 7.91% when the foragers foraged in an environment previously shaped by an intermediate or omniscient forager, respectively (middle and right boxplots in Figure 4). Similarly, the relative foraging efficiency of intermediate foragers compared to omniscient foragers increased of 10.31% and 9.62% when the forager foraged in an environment previously shaped by an intermediate or omniscient forager, respectively, compared to when the foragers foraged in an environment previously shaped by a naive forager. This means that space engineering has affected the foraging benefits of spatiotemporal memory, but not necessarily in a way that promotes the level of cognition that shaped it.

**Figure 4.**
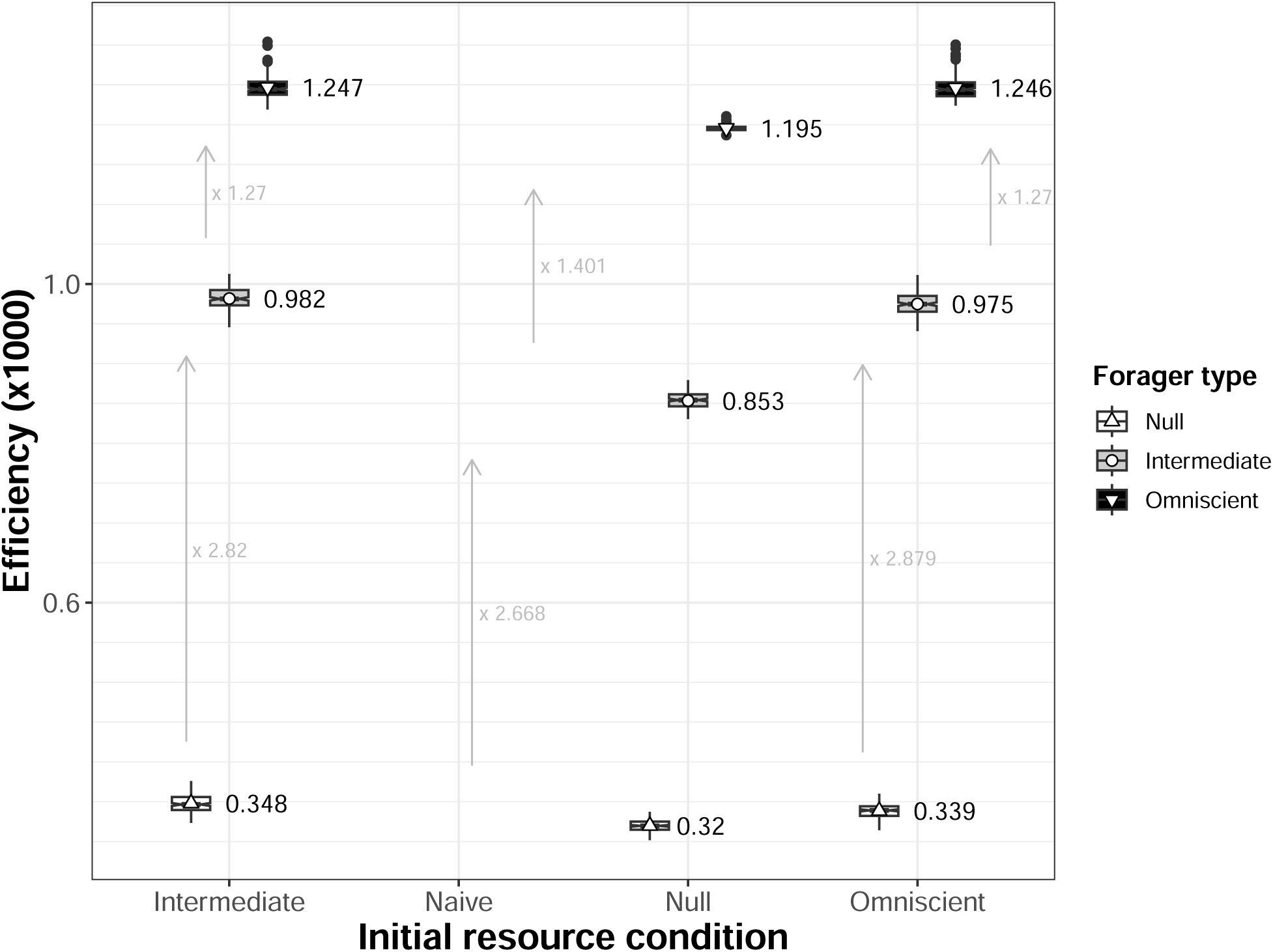
Relative foraging benefits of cognition are sensitive to the emerging resource distribution | The ‘initial condition’ of resource distribution was taken from the simulations in which a naive, intermediate or omniscient forager was first let foraging. Then, those three forager types were let foraging (separately) without dispersal and foraging efficiency was measured (cumulated food eaten over cumulated travelled distance).

### III.3 Route formation is reinforced because of opportunistic stops and by low plant space competition

How a forager moved to the target affected the emerging resource pattern (Figure 3B). In particular, moving to all plants encountered *en route* to the target (“All plants” moving rule, Figure 3B) induced a 1.117 times lower patchiness than only stopping to fruit plants (“Only fruiting plants” moving rule, Figure 3B) and 1.128 times lower than moving only to the target (“Only target plants” moving rule, Figure 3B). Resource alignment was 1.081 and 1.172 times higher when the forager stopped at all plants rather than only stopped fruiting plants or moved directly to the target (Figure 3B). Both patchiness and alignment were considerably reduced by competition for space between plants, with patchiness 1.643 and 2.134 times larger when competition for space between plants was low compared to when it was intermediate or high, and alignment being 2.04 and 7.814 times larger, respectively (Figure 3C). Spatial autocorrelation in fruiting date remained mostly unaffected by the forager movement rule but decreased with competition for space between plants with the same magnitude as the changes induced by the forager’s spatiotemporal knowledge (Figure 3B and C).

Variations in movement rules were paralleled by variations in routine behaviour, with foragers moving to all plants being 1.122 and 1.194 times more routine than foragers stopping only to fruit plants or their target, respectively. Competition for space between plants similarly affected the tendency of the forager to engage in routine behaviour, with a larger routine when competition for space between plants was low (Figure 3B).

## IV Discussion

**T**he environment is shaped by the species that inhabit it. The destructive behaviour of some animals when moving, such as forest elephants (Pringle, 2008; Vanleeuwe & Gautier-Hion,1998), can lead to blatant changes, but more elusive - yet no less ecologically relevant - engineering can also occur over longer time scales. Using an agent-based model, I have illustrated the role of vagile animals in shaping the distribution of plants through zoochory. Based on a minimal set of realistic assumptions, I showed how this slow process could affect resource distribution provided that the seed-dispersing animal was endowed with cognitive abilities, which in turn shaped selection on these abilities. Therefore, this study illustrates how foraging cognition could be both a cause and a consequence of resource heterogeneity. By its inherent simplicity, the models may nonetheless suffer from limitations which may weaken the conclusions in nature (see Box 1). As such, this model represents a proof of concept for an eco-evolutionary feedback loop between cognition and resource distribution, but does not confirm its existence in the wild.

As the breadth of knowledge increased, routine behaviour, the ability to repeatedly follow a fixed sequence of movements, developed (Riotte-Lambert et al., 2015). The model evidenced that the emergence of such behaviour was associated with changes in the distribution of resources, which tended to become more patchy, eventually forming linear features paralleling the routes used. The agglomeration of seeds for cognitive foragers has been proposed theoretically (John et al., 2016) and empirically (Fragoso et al., 2003), ultimately impacting the genetic structure and flow of plant populations (Gelmi-Candusso et al., 2017). Here, the model highlighted how engineering can go further by contributing to a more organised distribution of resources, with the emergence of linear features likely to be associated with the foraging routes used due to cognition, as suggested by forest elephant routes matching with fruit gradients in the forest (Blake et al., 2009). Indeed, the non-linearity of the changes in patchiness and alignment with the cognition of the forager likely corresponds to the formation of the routes themselves, with the “materialisation” first of the route intersections (hence higher patchiness, lower alignment for low cognition), where most of the recursions occur (Presotto et al., 2018), before more routes are used and formed when cognition increases. Long-distance travels through these routes may explain why animal-dispersed plant species are less aggregated than when other dispersal mechanisms are involved, such as barochory, where dispersed seeds do not travel far from the parent plant (Seidler & Plotkin, 2006). This engineering remained yet limited because the resource patterns did not fully match the simulated route-based patterns (see Appendix). Furthermore, given vertical inheritance of the parent plant phenology, seed dispersal induced a substantial shift in spatial autocorrelation in fruiting dates. This was not because the plants all ended up fruiting at the same time (i.e., due to global synchrony of plants), as there was still ample variation in fruiting timing at the end of the simulations (see Appendix). Plant synchrony is an important tool that can be used by foragers to plan their movements efficiently (Janmaat et al., 2012). Given the climatic drivers of large-scale synchrony, the latter can be tracked by animals (Janmaat et al., 2006). A fine-scale synchrony is also observed in nature (Lamontagne & Boutin, 2007) and might as likely affect animal foraging success. While large-scale population trends in phenology might stem from genetic constraints (Chuine et al., 2000; Lyngdoh et al., 2012), the emergence of spatial autocorrelation in fruiting date in this model (which considers vertical inheritance) reinforces this idea, and evidences how gene flow may affect, in addition to local abiotic factors (e.g., lightening, local temperature, etc.), the spatial contingency of phenological traits. However, these conclusions may be challenged by more accurate modelling of movement behaviour, which may lead to different results (see below). Despite these limits, the observed spatial engineering could still be sufficient for the emergence of eco-evolutionary feedback loops between animal resources and animal movement (and cognition) (Riotte-Lambert & Matthiopoulos, 2020) due to the evidenced repercussions on foragers’ foraging efficiency.

While the same individual might experience little change in resource distribution over its lifetime, the progressive engineering of resource distribution could overturn selection over generations. This model showed that changes in environmental conditions were likely to affect the benefits of memory, but not enough to completely overturn selection. Indeed, in all environment, memory was advantageous. The relative advantage of omniscience compared to no memory was nearly constant whatever the environment. Interestingly in contrast, the relative advantage of intermediate memory compare to no memory, was increased in engineered environments. The performance of these naive agents is clearly dependent on their sensory abilities (Grove, 2013). Although always superior to the randomly expected nearest neighbour distance, the sensory range modelled here may have been insufficient for simple movement strategies such as taxis/kinesis to perform well. As much as memory could be favoured, the environment could also shape sensory abilities (LaScala-Gruenewald et al., 2019). Still, the changes in the relative efficiency of cognitive agents open the door to the possible emergence of evolutionary feedback loops if the costs of memory are no longer compensated, which should lead to a dynamic equilibrium (Beisner et al., 2003) due to the simultaneous changes in memory benefits (hence selection) with space engineering.

Space engineering was reinforced by the conjunction of two forces: the forager movement rules and competition for space between plants. Maximum resource engineering was achieved when the individual moved to all the plants it encountered along the way and was able to learn the new seedling locations (see Sensitivity to the learning pathways following old plant mortality). At the other extreme, when the animal moved directly to the best target, and only learned previously visited trees in case an old tree died, engineering was reduced, following routine trends. Although more efficient, moving straight to the target is rarely observed in nature. Instead, animals tend to move along routes, sometimes bypassing or otherwise opportunistically inspecting plants without resources, fitting with a beacon-based navigation, where individuals move from beacon to beacon to reach a goal (Fagan et al., 2013; Warren, 2019). Such a movement strategy gives them the possibility to monitor the environment (de Guinea et al., 2019), allowing them to update personal information. As such, despite short-term costs due to detours, moving to all trees might provide long-term advantages. However, the simulated movement patterns remain extremely simplistic. Animal movement, learning, and memory, is in reality much more nuanced than in this modelling work (see Box 1). This could be important for seed acquisition and deposition (Morales & Morán López, 2022). For example, this model considered directed movement supported by memory, which could alternate with random steps when memory was limited but independent of local cue feedback, as it would rather be expected for efficient foraging (Benhamou & Collet, 2015). Animal movement may indeed alternate between ‘large-scale’ directed relocation events and ‘area-restricted’ wandering searches of the plant in a non-random manner (Bartumeus et al., 2016). When and where area-restricted searches occur may further contribute to the emergent distribution of seed deposition (Russo et al., 2006b; Westcott et al., 2005), provided that both seed deposition and foraging can occur simultaneously. Animals may yet be selective about where and when they defecate. The presence of latrines may contribute to shaping the distribution of resources too (Fragoso, 1997; Fuzessy et al., 2022b; Irwin et al., 2004; Pouvelle et al., 2009; Sakamoto & Takatsuki, 2015). Overall, as movement and/or dispersal scales with behaviour and personality (Brehm & Mortelliti, 2022; Zwolak, 2018; Zwolak & Sih, 2020), but also species and individual size (Jordano et al., 2007), a multi-individual and multi-species system could lead to a more complex multi-scale scaffolding than illustrated here (discussed in Box 1). This could explain why the evolution of the cognitive machinery underpinning foraging might depend on the frugivore community composition (Robira & Perez-Lamarque, 2023).

In this scaffolding, the plant itself may also play a role. Not only are seed characteristics important in the dispersal process (Albert et al., 2015; Fuzessy et al., 2018), but the success of seedling germination is highly dependent on the conditions in which the seed is deposited, due to direct or indirect actions of conspecifics. This model took into account negative interactions between plants (e.g., competition for resource, Craine & Dybzinski, 2013, inducing an exclusive space use) and illustrated how this could act as a barrier to spatial engineering. Indeed, as competition for space between plants increased, both patchiness and alignment logically decreased, cancelling out the engineering effect of cognition. While the model effect was interpreted as a competition for space, this may also encompass other processes such as greater exposure of seeds to predation risk in the vicinity of adult plant conspecifics (Janzen, 1970). Conversely, facilitation could also occur, with the adult plants acting as a shelter for new seedlings [Callaway (1995); but see Connell (1971) and Janzen (1970) for a contrasting rationale]. This facilitation should increase clustering. While directly unmodelled here, in the quasi absence of competition for space, plant range was coherently drastically reduced (i.e., shrunk; Figure A5), with a tendency to agglomerate towards the centre of the map, where individual recurrences are likely to be most important. Therefore, the engineered patterns (e.g., from increased patchiness to route formation) may also depend on plant interactions, adding to the complexity of the system depicted.

The sessile lifestyle of plants may have created the illusion of residual fixity in the environmental conditions that shape the lives of frugivorous animals. While, in the absence of depletion, the resource spatial distribution is likely to remain the same in the short term, conferring the benefits of information retention in frugivores, resource spatial distribution can also change over longer periods of time. Selection could thus operate in two ways: frugivores could play a key role in plant traits by actively selecting and dispersing their food, while their traits could have been shaped by foraging on that primary food. Considering the feedback loops between these different entities could be an important facet to make sense of still poorly understood patterns of co-evolution, such as the primates/angiosperms dyad (Fuzessy et al., 2023; Fuzessy et al., 2022a; Gómez & Verdú, 2012; Sussman, 1991).

#### Box 1 Taking a step back: a critical discussion of the model limitations

This model is a proof of concept (i.e., answers the “what if” question, Morales & Moran Lopez, 2022): it shows that an eco-evolutionary feedback loop between animal cognition and resource distribution is possible, but it does not confirm its existence in nature. This is because, paradoxically, this model represents both a field of possibilities much larger than its realisation in nature, and at the same time a drastic simplification of reality, which is actually the essence of the model’s interest (Morales & Moran Lopez, 2022, as is often said, “all models are wrong, but some are more useful than others”, Georges E.P. Box). After some abstraction, this simplification makes it possible to find analogies with various natural systems and make predictions about what might happen in the wild. This is partly due to the arbitrary choice of spatial and temporal units, which allows the model to be transferred to different ecological systems. If the temporal unit is a day and the spatial unit is a metre, and spatial competition is low, a tropical rainforest with asynchronous plants can be considered (e.g., many of the fig tree species used by animals as a staple food, Shanahan et al., 2001). If you change the time unit to half a day (and thus reduce the speed by two to keep the absolute speed the same), plants can be considered synchronous, and thus you might consider focusing on the fruiting season only of a seasonal rainforest, that classically characterises some rainforests of Central Africa and can influence animals’ feeding habits and movement (Masi et al., 2009; Terborgh, 1986). However, the mechanistic simplification of a real system also imposes limits on the generalisability of the model, calling into question the accuracy of the predictions. This box highlights and discusses some of the model limitations, and can serve as a compass for future research.

#### The environment: a shared place

The simulations presented here focused on a single forager occupying a square area filled with plants. While this postulate may be valid for a territorial species (Burt, 1943), or because of cognitively induced passive spatial segregation (Riotte-Lambert et al., 2015), some species may still have overlap with conspecifics (e.g., in primates: Pearce et al., 2013). Moreover, space is shared not only with individuals of the same species, but also with other species, some of which may be targeting similar resources. Because individuals compete or learn from other individuals, they may thus alter their movement based on conspecific (Sassi et al., in press) or heterospecific (Havmøller et al., 2021) movements. Furthermore, the direct or indirect interaction between individuals is likely to shape the selection on their cognition too (Ashton et al., 2020; Robira & Perez-Lamarque, 2023). As individuals with different levels of cognitive types can coexist, they can influence how animals move and deposit seeds (Borah & Beckman, 2022; Zwolak & Sih, 2020). By allowing different individuals to forage, and thus for the possible coexistence of “opposite” movement patterns, the conclusions presented here may well be overturned. Therefore, multi-individual models should be a future focus to understand the role of the community (rather than the individual) in eco-evolutionary feedback loops between cognition and resource distribution.

#### The coexistence of dispersal syndromes

The only way for the modelled plants to disperse was by zoochory. The amalgamation of a diaspore and a dispersal mode (i.e., the existence of a dispersal syndrome) is common, but incorrect (Green et al., 2022). Many of the fleshy fruits, for example, are dispersed by both barochory (local dispersal) and zoochory (potentially “global” dispersal). The multi-channel possibilities for dispersal are likely to affect the resulting patterns of seed deposition (Clark et al., 1999; Rogers et al., 2019). By considering zoochory only, this model implicitly assumes that it is the dominating channel of dispersal and therefore, that space engineering may only be substantial, and affect selection on cognition, in that case. To which extent barochory (or other dispersal means) may cancel out the engineering highlighted here, and thus, the consequence on animals’ foraging cognition, remains to be explored.

#### Movement as a multifaceted behavioural trait

Animals may not move at random: their movement might be planned and result in a succession of decisions inducing different movement modes (Fryxell et al., 2008). For example, an animal may move from a locally exploited patch to another patch (relocation) and then search for the resource within that patch (exploitation phase, Bartumeus et al., 2016; Benhamou & Collet, 2015). Animal decisions, and thus, the multiplicity of movement modes, may result from changes in both animal condition and environmental conditions (Nathan et al., 2008). For example, during periods of food scarcity, hungry animals can drastically reduce their movement, for example by entering torpor (Vuarin & Henry, 2014), as it has been modelled here. In contrast, satiated individuals may engage less in foraging movement (Hansen et al., 2015). Such short-term changes, following food distribution and animal physiology, may even persist over longer periods of time. For example, western gorillas prefer fruit to leaves. However, the abundance of fruit varies throughout the year, as does their diet, which changes from high to low frugivory (Remis, 1997). These variations are mimicked by oscillations in their activity, the distance they travel, and the area they cover per day (Doran-Sheehy et al., 2004; Masi et al., 2009; Robira et al., 2023b). None of these possibilities occurred in this model. Agent movement was restricted to the simplest case: ballistic movement from target plant to target plant, if known and of interest, throughout the season. Only variations of opportunism were implemented: stopping at all or only fruiting plants encountered *en route*, or none. This was shown to reinforce, or buffer, the spatial engineering following the seed dispersal by the agents. It is therefore likely that the neglect of movement pattern diversity, and in particular concentrated movement such as area restricted search, may have minimised long-term resource aggregation. Overall, coupling agent-based models such as this with more explicit behaviourally-dependent habitat use and movement patterns (Bialozyt et al., 2014; Russo et al., 2006a) would increase the predictive power of these modelling works.

For eco-evolutionary feedback loops to occur, ecological changes (resulting from agents’ actions) should be substantial, and these ecological changes should in turn induce changes in agents’ traits (Post & Palkovacs, 2009). This model formally illustrates the first condition (Q1; see the third next bullet point) and suggests that the second condition is true, as the benefits of spatiotemporal memory vary with the resulting engineering of resource distribution (Q2). It only suggests this because agents’ fitness is only proxied by foraging efficiency (see the second next bullet point). Furthermore, this model excludes the evolution of the agents (spatio-temporal memory width does not change during the simulation). This is particularly important in explaining why the model was designed to accelerate ecological change (e.g., plants reaching maturity quickly after dispersal) in order to limit the computational time needed to test the first condition (for substantial ecological change), but these choices should be inconsequential on the model conclusion given the current model design (i.e., absence of evolution). However, in eco-evolutionary feedbacks, the rate at which the environment changes and the species traits of interest evolve (here spatio-temporal memory) are key. Indeed, for eco-evolutionary feedbacks to arise and be maintained, the ecological and evolutionary time scales must be congruent (Post & Palkovacs, 2009). Here, only ecological changes occurred, evolution did not occur in parallel. To unambiguously confirm the existence of eco-evo feedbacks, explicit modelling of spatio-temporal evolution should be performed. This would require careful consideration of the ecological processes underlying seed dispersal success and plant growth patterns. It would also mean considering a mortal forager. Indeed, in the present simulations, the forager could be considered immortal or simply replaced by a clone with perfect vertical inheritance of knowledge, a rather unrealistic assumption. Moreover, an individual may also show plasticity in the known information (see the next bullet point below). Considering learning and inheritance of knowledge is an essential next step to explicitly model the evolution of spatiotemporal memory and thus to challenge the conclusions of this model.

#### Learning and forgetting: the dual aspect of memory

In this model, the agent was rarely able to learn new profitable locations. Only when a known plant died was the location of an unknown plant (either the dispersed seed in most simulations, or the most recently visited fruiting plant, see Appendix, Sensitivity to the learning pathways following old plant mortality) learned. Such a simplified learning mechanism, which is clearly unrealistic (especially for instantaneous learning of instantaneously growing new seeds), was a convenient way to keep plant density (hence instantaneous growth) and memory size (hence instantaneous learning) constant, as these two aspects are crucial in determining the benefits of memory (Boyer & Walsh, 2010; Bracis et al., 2015; Grove, 2013, this paper). However, the consequence of memory may arise from the nexus of learning, knowing, and forgetting (Kraemer & Golding, 1997; Tello-Ramos et al., 2019). In particular, in the case of foragers’ recursions, the speed with which a fruitful site can be memorised or erased, and the cognitive abilities and dynamics in general, may have a significant impact on foragers’ patterns of revisits and routine (Berger-Tal & Bar-David, 2015), both of which were prerequisites for the emergence of resource aggregation and local correlation in phenology. This would explain why the two learning pathways produced qualitatively identical (i.e., ranging similarly compared to the benchmarks), yet quantitatively unidentical results, with memorisation of the seedling accentuating the environmental engineering effect. Relaxing assumptions on memory stability and learning speed would necessarily affect further the speed at which e2c2ological change may occur, as well as the likelihood for memory to evolve. Further work should therefore consider a more dynamic memory to see how this may affect the existence and nature of eco-evolutionary feedback loops involved in memory evolution.

#### The memory burden: memory benefits does not necessarily imply selection

For a trait to be advantageous, not only must it confer some benefits, but these benefits must outweigh the costs that the trait may impose. In the case of cognition, for example, it is known that it imposes additional metabolic costs, while less straightforward costs may arise from the need to learn, from memory impairment, or from the consequences of knowledge inaccuracy (Burns et al., 2011; Fagan et al., 2013). These costs are rarely studied and quantified empirically, highlighting the need for such research to make models such as this more realistic and less condemned to speculation about memory adaptiveness. In the meantime, in the absence of explicit costs, the shortcut taken here has been to assume that foraging benefits are synonymous with fitness benefits. In other words, the benefits of foraging compensate for the additional costs of cognition. However, this is not necessarily the case. For example, the benefits of spatiotemporal memory vary significantly with patterns of spatial distribution of resources (Grove, 2013) and phenology (Robira et al., 2021). Variation in benefits with environment conditions explains why cognition may vary between species (Van Woerden et al., 2010; van Woerden et al., 2012). A clear formulation of fitness, and hence of the benefit-cost trade-off associated with memory, is essential to further elaborate on the role of cognition in animal evolution, including through these (possible) eco-evolutionary feedback loops (Hoyle et al., 2008).

#### Biological substantiality: what is a lot in ecology and evolution?

As should be the case in modelling work, differences in resource patterns have been estimated by quantifying changes in some indices compared to expected changes estimated on the basis of simulated data, rather than on the basis of statistical tests which are meaningless in this case (White et al., 2014). However, while this characterises the ecological relevance of the change, it does not characterise the eco-evolutionary relevance of the consequence of the change. If that consequence (e.g., effects on foraging benefits or fitness) is a nonlinear function of the ecological variable of interest (e.g., patchiness, alignment, spatial autocorrelation, etc.), ridiculously small or large differences in this variable compared to the true range of possibilities, may perhaps have in the end very large, or on the contrary very low, evolutionary consequence. This was the reason behind directly measuring the changes in foraging efficiency of foragers following the changes in resource spatio-temporal distribution (Q2). However, foraging efficiency is only an indirect proxy for individual fitness: not only because it does not measure costs (see above), but also because the functional form linking foraging efficiency to fitness is unknown. While this approximation is common in most, if not all, empirical or theoretical studies on cognitive foraging, it drastically limits the predictive power of this work specifically: while results put forward that an eco-evolutionary feedback loop between cognition and resources may exist, there is no certainty that it does so in nature, and that this in itself has any consequences for the evolutionary trajectories of both foragers and plants. This is also why this model does not intend to portray a fully realistic situation. However, by demonstrating that an eco-evolutionary feedback loop between animal cognition and resource distribution can be relevant under a minimal set of assumptions, it now justifies the undertaking of empirical studies to test it *in vivo*.

## V Acknowledgements

Simulations were run on the HPC cluster of the Edmund Mach Foundation. I thank Louise Riotte-Lambert and Kavel Ozturk for discussion on the model rationale and feedback on the manuscript, Eva Gril for discussion on botanical assumptions, as well as Alexandra Loveridge for English editing. I thank Fabien Laroche (PCI recommender), Julia Astegiano, Christophe Baltzinger and another anonymous reviewer for feedback on a previous version of the manuscript.

## VI Funding

No specific funding was obtained for this research.

## VII Declaration of conflict of interest

I declare having no conflict of interest.

## VIII Data and code availability

The code used to generate the data and perform the analyses is available at https://github.com/benjaminrobira/ModelZoochoryCognition. To fully reproduce the analyses, a singularity image (through which the model can be run and which was used to run the simulations on the HPC cluster) can be provided on request, as it is too large to store on git.

## IX Appendix

### IX.1 Using patchiness and alignment to characterise spatial point patterns

I provide below evidence that the two metrics, namely *patchiness* and *alignment*, allows for distinguishing between homogeneous, heterogeneous and route point patterns. In both the low (100 plants) and high (1000 plants) density, I considered 1/10 of the number of plants as the number of clusters for the heterogeneous distribution (i.e., 10 clusters at low density, 100 at high density). Around these clusters of strictly equivalent size, plants were distributed according to a Gaussian distribution (both for the *x* and *y* axis) of variance 50 su. The distribution resembling a route network was obtained by modelling one (low density) to ten (high density) correlated random walks of 100 steps (“simm.crw” function of the *adehabitatLT* package, Calenge et al. (2023), with a scaling parameter, to determine step length, of 20 su, and a concentration parameter, relative to the correlation in heading, of 0.9). I calculated the two metrics on 200 distributions.

Homogeneous distributions are identified by a low patchiness and a low alignment. Heterogeneous distributions are identified by an intermediate patchiness and a low alignment. Finally route point distributions are evidenced by a high patchiness and a high alignment (Figure A2). This pattern is robust to density variations, although absolute differences in the indices shrink with density increase (Figure A2). The benchmark values at high plant density (1000 plants) are are depicted in Figure 3 of the main text.

### IX.2 Using Moran’s Index to characterise plant synchrony

Below I provide evidence that Moran’s Index, adapted for circular variables (see main text), can correctly identify environments with high or low local synchrony in fruiting dates. To do this, I simulated square environments of length 1000 su and containing 1000 plants either homogeneously or heterogeneously distributed (as above, Figure A3). I assigned each plant a start of fruiting date (in radians) to mimic an environment with no, intermediate or local synchrony (Figure A3). To do this, I first initiated the fruiting dates for 50 plants by randomly sampling in a wrapped Gaussian distribution with a concentration parameter equal to 0.5 and centred on 0 (“rwrappednormal” function of the *circular* package Agostinelli & Lund, 2023). Then, for the 950 remaining plants, I sampled their fruiting dates in a wrapped Gaussian distribution with a concentration parameter equal to 0 (low local synchrony), 0.5 (medium local synchrony), or 1 (high local synchrony), centred on the fruiting date of the nearest plant (with a fruiting date already associated). Then, to smooth the pattern in space, I corrected the fruiting dates of all plants by taking the weighted (by the inverse of the Euclidean distance; the value of the weight for the focal plant was set to 1) circular mean (“weighted.mean.circular” function of the *circular* package Agostinelli & Lund, 2023), considering only the 50 closest plants (including the focal plant itself). I simulated 200 environments for each spatial (homogeneous or heterogeneous) and temporal (i.e., synchrony level) pattern. Overall, Moran’s Index increased coherently with simulated synchrony, regardless of spatial distribution (Figure A3). The spatial distribution affected the resulting Moran’s Index, with a spatial autocorrelation 1.38 times superior for a simulated same synchrony when the environment was heterogeneous compared to homogeneous. The benchmark values are depicted in Figure 3 of the main text.

**Figure A1.**
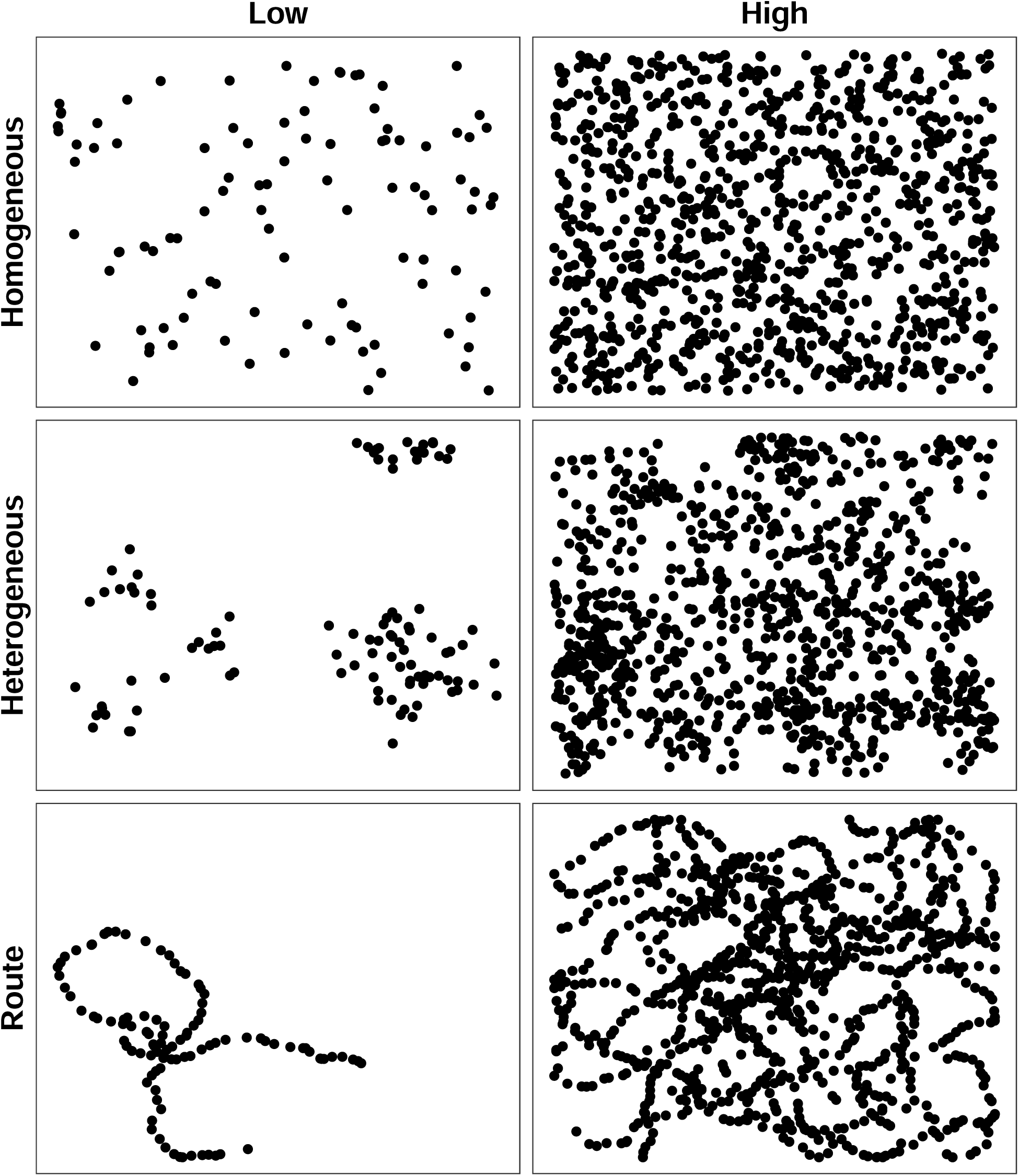
Simulated spatial distributions to investigate metrics value | Distribution: Homo. = homogeneous, Hetero. = Heterogeneous, Route = Route network-like. plant population size: Low = 100 plants, High = 1000 plants, within a square map of side length 1000 su.

**Figure A2.**
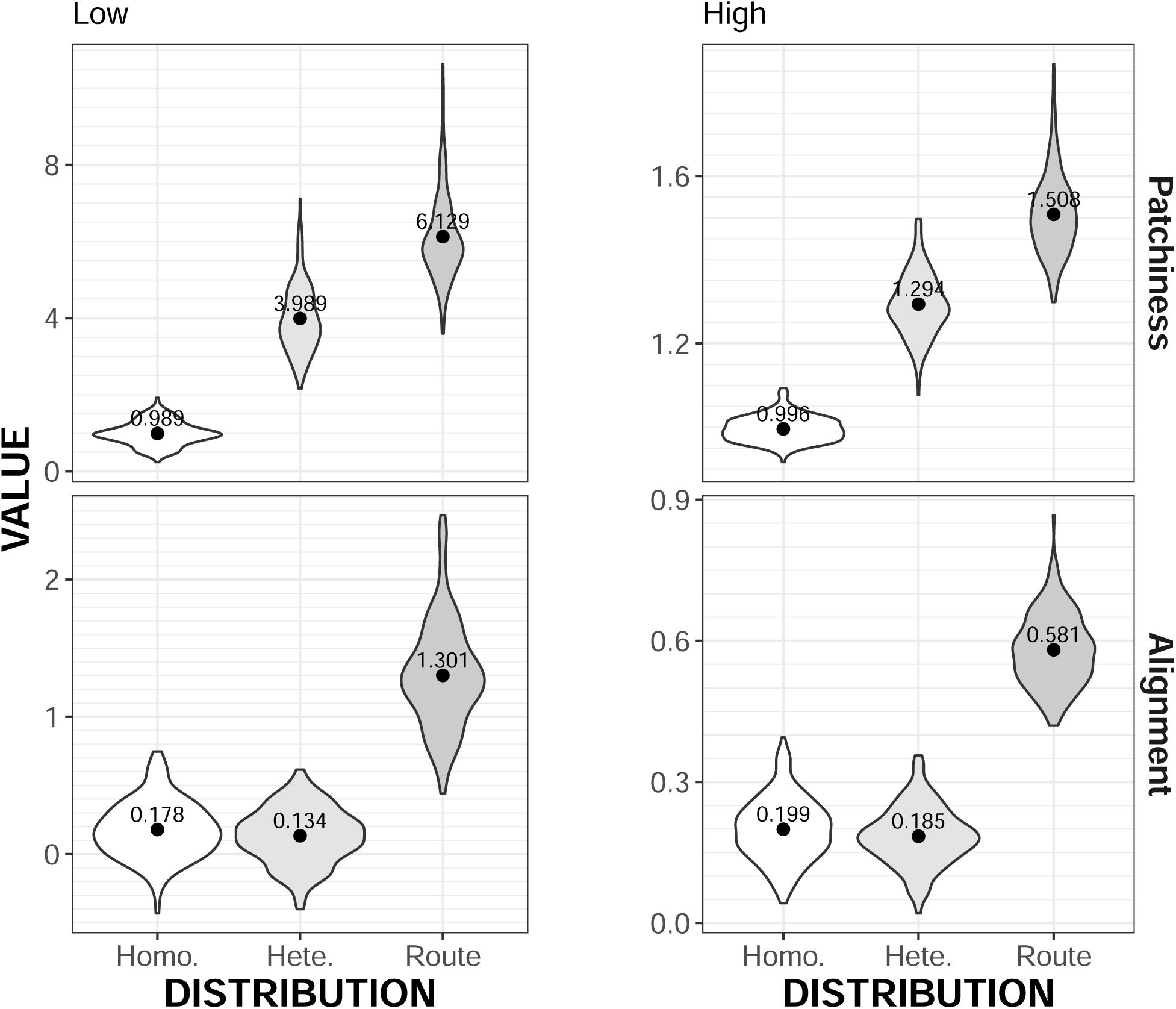
Patchiness and alignment allow for identifying spatial distributions at low (100 plants) or high (1000 plants) plant population size | Results are based from simulated data following a homogeneous (Homo.), heterogeneous (Hetero.) or route network-like (Route) distribution. The mean value is depicted by the black dot and the text. The shape of the distribution is depicted by the contour of the violins.

**Figure A3.**
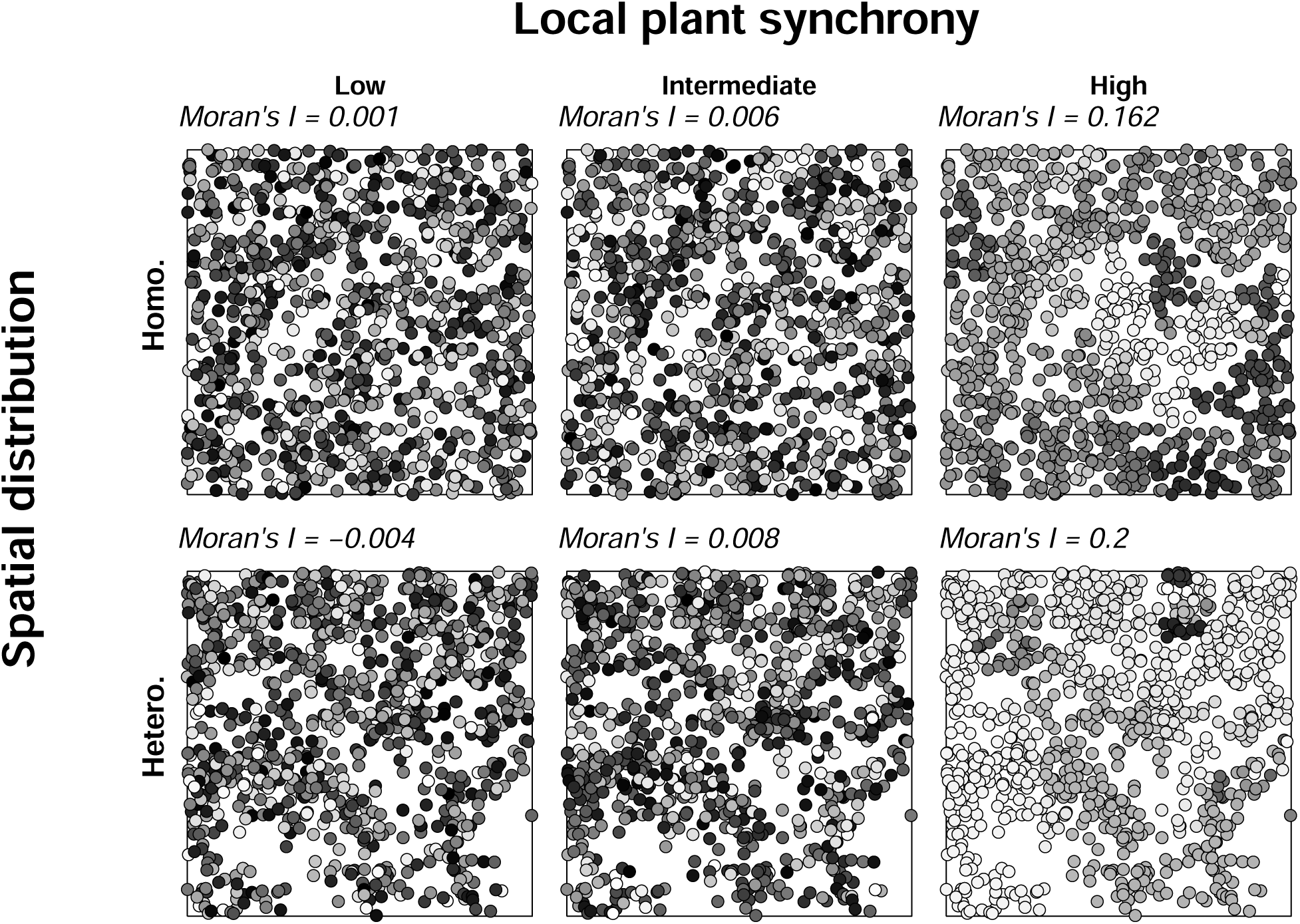
Examples of simulated environments varying in their spatial and temporal distribution | The dots represent the plants. They are coloured along a white-to-black gradient according to the value of their fruiting date. The more synchronous two plants are, the more their colours match.

**Figure A4.**
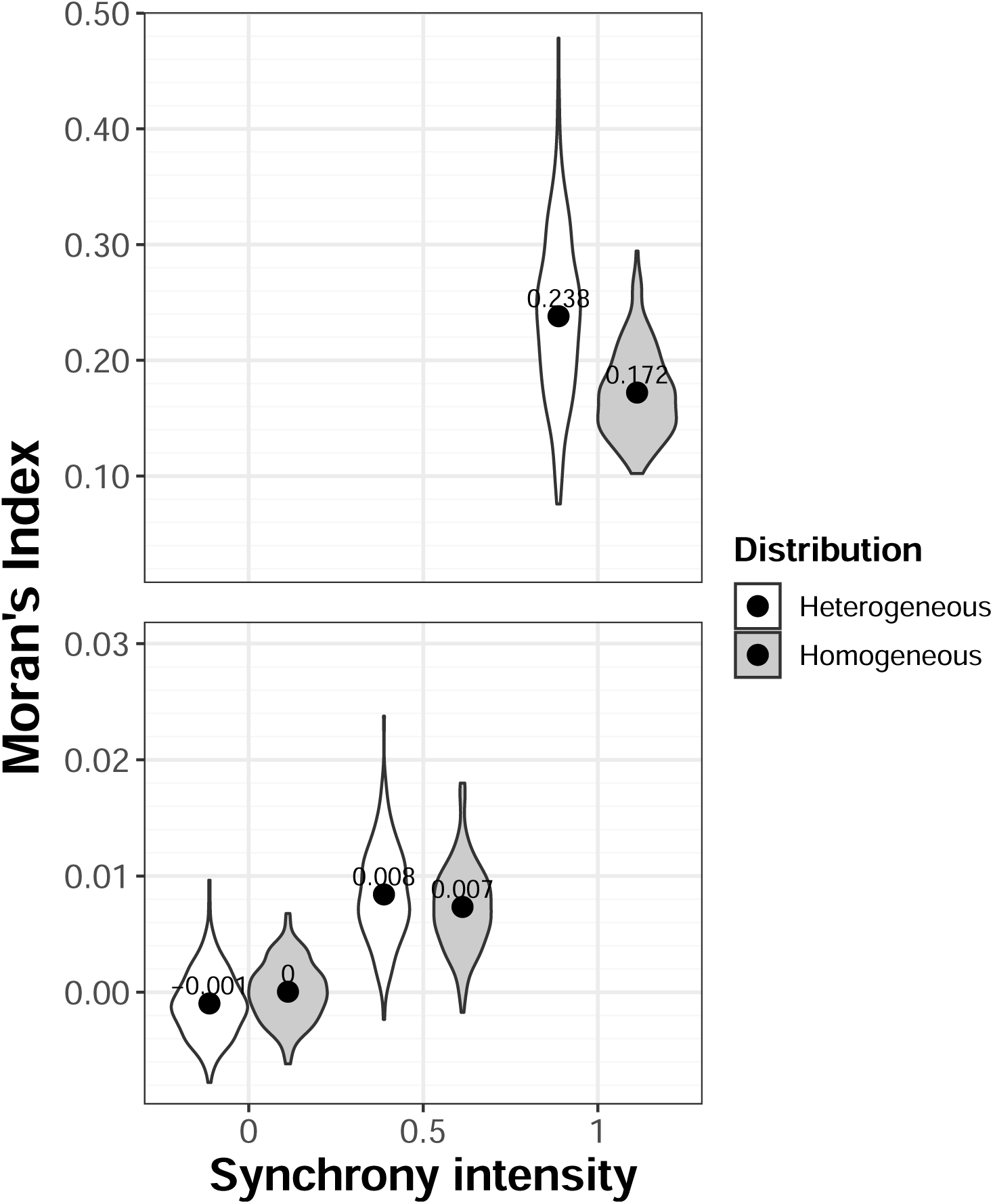
Moran’s Index adapted for circular variables correctly identifies local synchrony among plants | Results are based from simulated data following a homogeneous (Homo.) or heterogeneous (Hetero.) spatial distribution. The synchrony intensity is related to the concentration parameter used when sampling in a wrapped Gaussian distribution (low = 0, intermediate = 0.5, high = 1). The mean value of Moran’s Index is depicted by the black dot and the text. The shape of the distribution is depicted by the contour of the violins. Note the non-linearity on the *y* axis. This is used to highlight the differences between low and intermediate synchrony.

### IX.3 Variations in resource range shrinkage intensity

Shrinkage was affected by the forager’s cognition, peaking at low cognition (Figure A5a). Shrinkage was also driven by the forager limiting visits to plant encountered *en route*, with a maximum reached when the forager discarded all of these plants and moved directly to the target (Figure A5b). Finally, shrinkage was removed when the competition for space between plants was high (Figure A5c).

**Figure A5.**
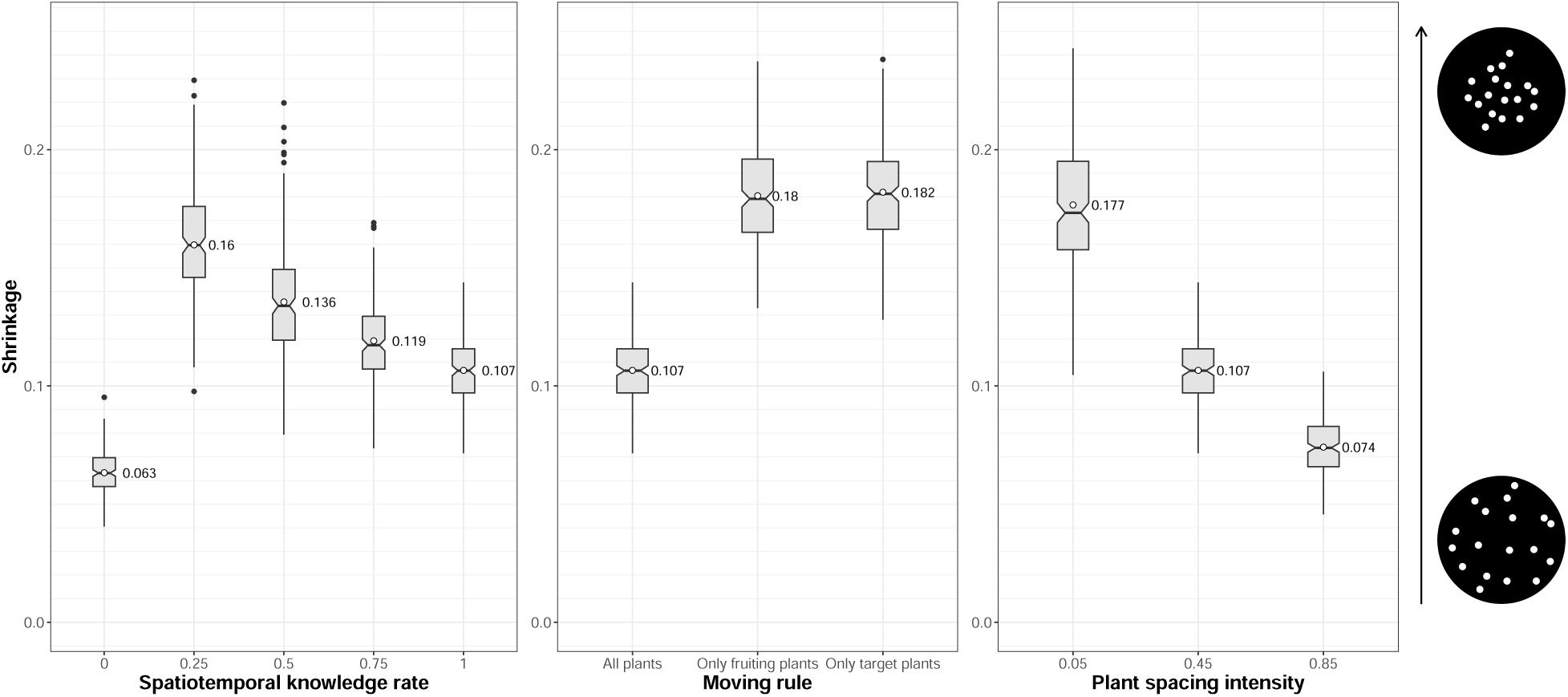
Shrinkage of resource range is driven by cognition, the forager’s movement rules and competition for space between plants.

### IX.4 Variations in routine behaviour when no dispersal occurs

In the absence of seed dispersal (scenarios in Q2 were used), routine behaviour still increased (and saturated) with more advanced cognition, whatever the environment (i.e., the engineering level, Figure A6).

### IX.5 Variations in fruiting dates

Considering the fruiting dates as circular variables (i.e., vectors with a length of one unit and a given direction), the distribution of fruiting dates can be summarised by a mean vector, the direction of which characterises the mean fruiting date and the length of which characterises the variance around this mean (with a length of 1 characterising low variance, Batschelet, 1981). I thus quantified the variation in fruiting dates as one minus the length of the mean vector of fruiting dates (Batschelet, 1981) in either the initial or final conditions. It varies from 0 (no variation; all fruiting dates are the same) to 1 (extensive variation; fruiting dates are uniformly distributed). At the beginning or end of the simulations, the variation in fruiting dates was considerable, regardless of the scenario tested (Figure A7).

**Figure A6.**
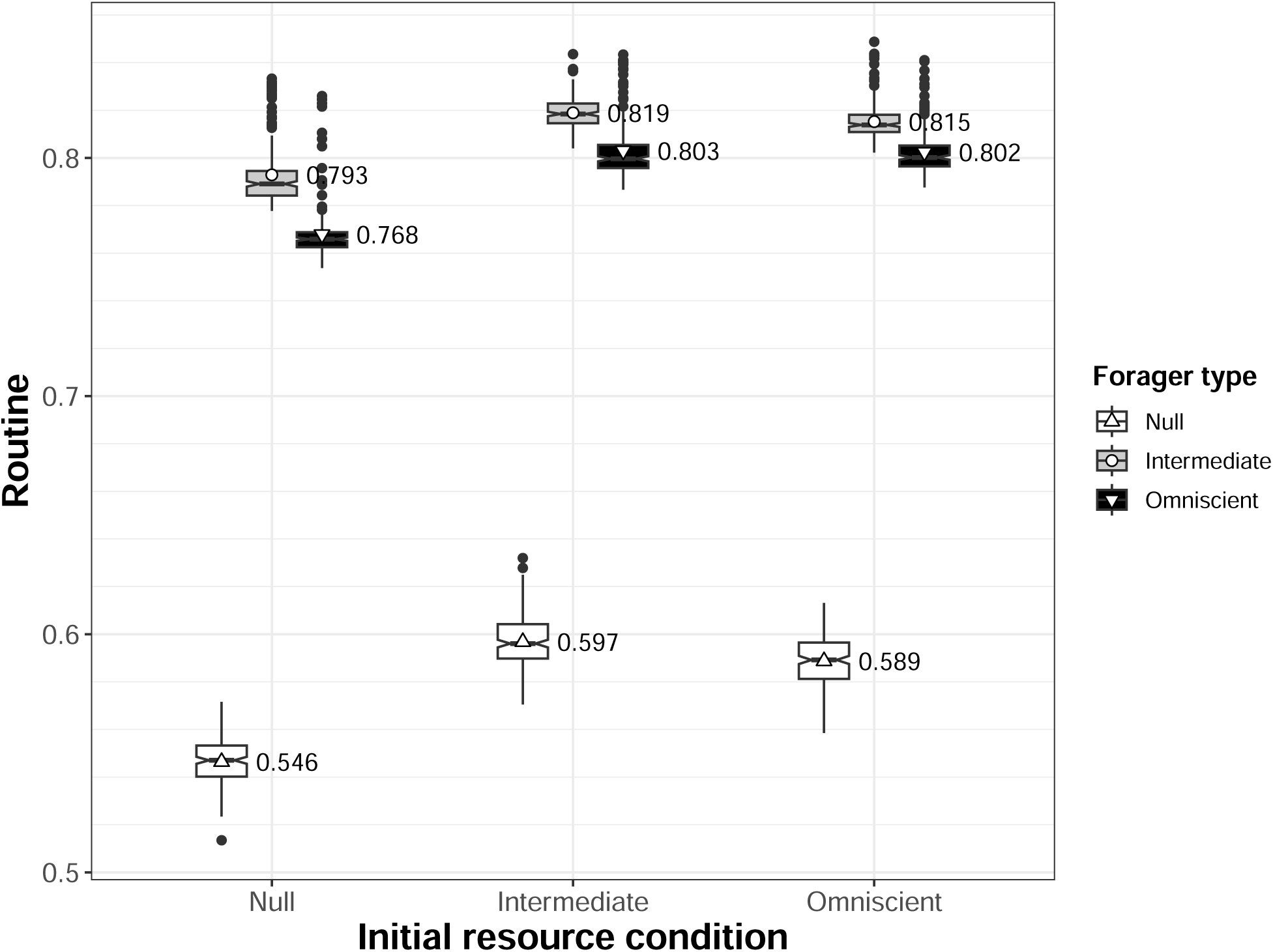
Routine emerges when cognition is high, independent of environment engineering | The ‘initial condition’ of resource distribution was taken from the simulations in which a naive, intermediate or omniscient forager was first let foraging. Then, those three forager types were let foraging (separately) without dispersal and routine behaviour was quantified (the tendency of the forager to target plants in a predictible way; values close to 0 indicate no predictible movements, values close to 1 indicate highly predictible movements).

**Figure A7.**
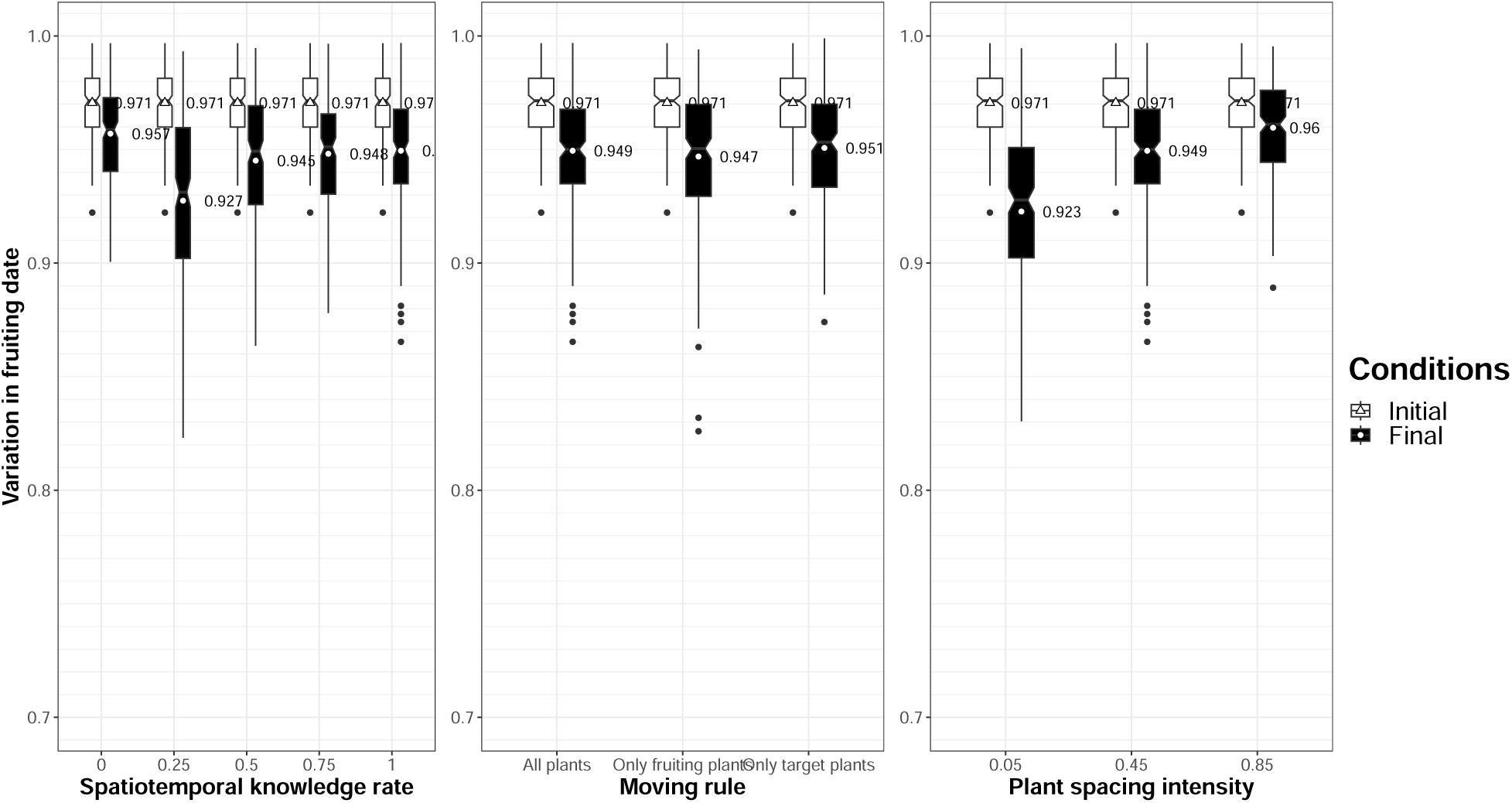
Variation in fruiting dates at the beginning of the simulation (‘initial’ condition) or at the end (‘final’ condition) | The plots show the results of 200 simulations when a seed-dispersing forager with varying level of cognition (i.e., proportion of plant locations and associated phenology known), moving choice rules or space competition intensities between plants (the two latter being simulated with an omniscient forager; left to right) was let foraging for a long time. The index of variation in fruiting dates (one minus the length of the mean vector of fruiting dates, Batschelet, 1981) ranges from 0 (no variation) to 1 (extensive variation).

### IX.6 Sensitivity to the agent’s speed

The emergence of spatial autocorrelation in fruiting dates depends on how far the seed is from the parent plant. Therefore, it may depend not only on the routine of the agent, but also on the speed at which it moves, and thus on the area that an agent can explore during the dispersal window. To investigate the robustness of the observation to the agent’s speed of movement (and thus allow analogies with other biological systems, since it is this speed that establishes the link between the ‘arbitrary’ spatial and temporal units), I ran additional simulations considering an omniscient forager moving at either half (‘low speed’) or twice (‘high speed’) the speed used for the standard simulations in the main text (‘intermediate speed’).

In comparison to other forager’s traits and in the range tested, movement speed was not the main driver of the resource spatiotemporal distribution (Figure A8A).

### IX.7 Sensitivity to the learning pathways following old plant mortality

In the model, when the deposited seed was able to grow, it caused the death of another random adult plant. Since memory size is an important feature that determines the benefits of memory in foraging (Figure 4), the new seed location and phenology was immediately known to the agent when a known plant died. To investigate whether this had undesirable consequences for the model’s results, I simulated the behaviour of an agent endowed with cognitive but not perfect knowledge (spatiotemporal knowledge rate of 0.25, 0.5, 0.75) learning the location and phenology of the most recently seen fruiting plant when another adult plant died (‘Recent fruiting plant learned’) and compared the simulation results (patchiness, alignment, spatial autocorrelation and routine) with those presented in the main text when the seedling location was learned (‘Seedling learned’).

The inability of the agent to directly learn the seedling location and phenology when an old plant died, and only the location and phenology of the most recently visited fruiting plant, limited the level of engineering (lower patchiness, alignment and spatial autocorrelation in fruiting dates, Figure A8B). However, both learning pathways ranged similarly compared to the benchmarks, so the overall pattern of engineering remained largely the same (Figure A8B).

**Figure A8.**
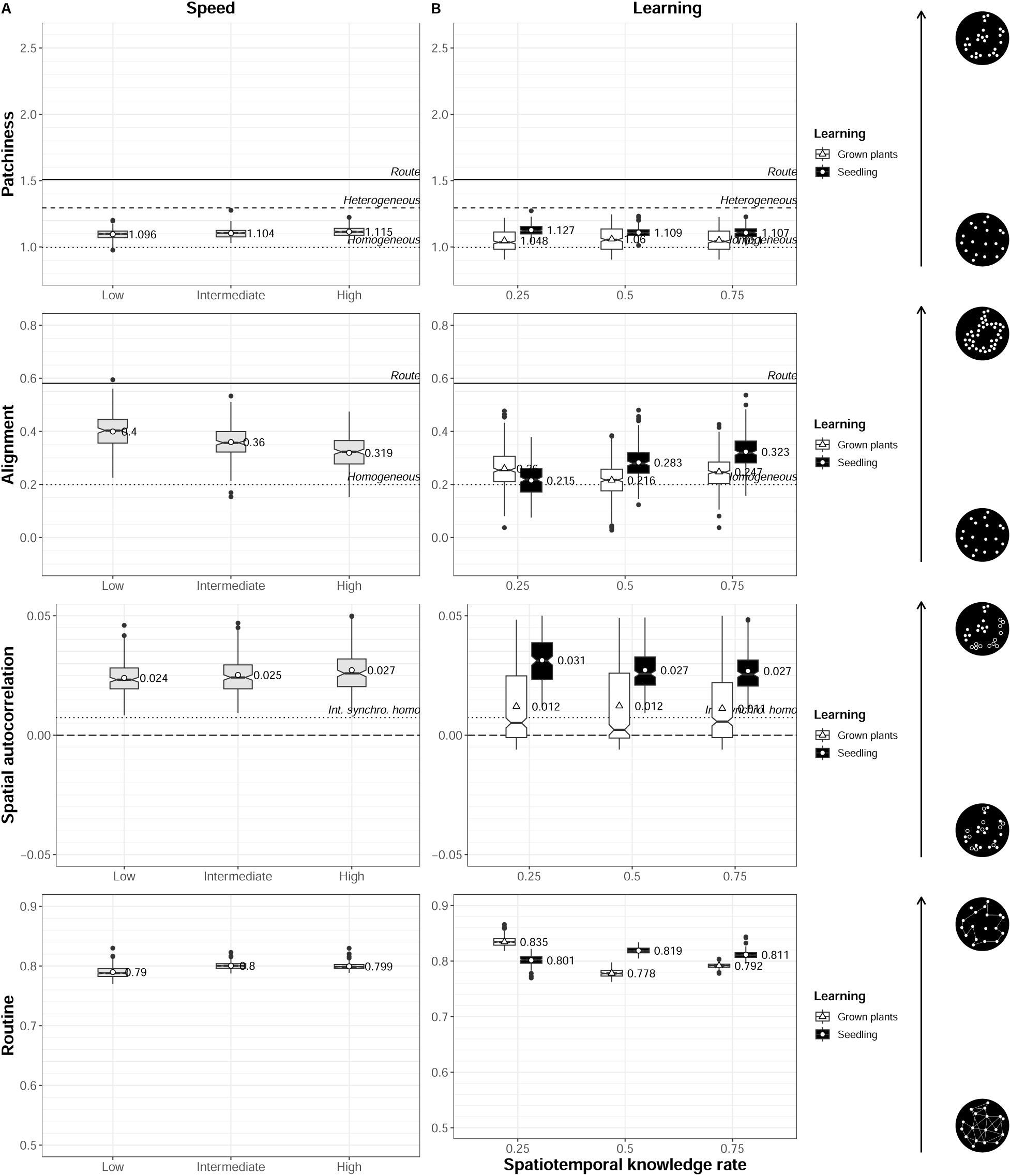
Seed-dispersing foragers can shape resource distribution | The notched box plots show the results of 200 simulations when a seed-dispersing forager with varying speed or learning abilities was let foraging for a long time. The resulting distribution pattern was assessed by its patchiness (the tendency of plants to be aggregated into patches; larger values indicate highly aggregated plans), its alignment (the tendency of plants to form linear geometries; larger values indicate the presence of linear geometries) and spatial {autocorrelation} in fruiting date (the tendency of plants to have fruiting dates closer/farther in time to their spatial neighbours; values close to -1 indicate negative spatial autocorrelation, values close to 0 indicate no spatial autocorrelation, values close to 1 indicate positive spatial autocorrelation). Indices’ benchmarks obtained from simulated environments are represented by the horizontal lines (Homo. = Homogeneous, Hetero. = Heterogeneous, Synchro. = Synchrony) (see Appendix). Because of overlap, only alignment and intermediate synchrony in a homogeneous (and not heterogeneous) environment was plotted. The emerging movement behaviour of the forager was described in terms of routine behaviour (the tendency of the forager to target plans in a predictible way; values close to 0 indicate no predictible movements, values close to 1 indicate highly predictible movements).

## Notes

### Competing Interest Statement

The authors have declared no competing interest.

### Summary of Updates

Manuscript revised after reviewers' feedback: added simulations, added discussion, overall text improvement, added figures.

## References

Addis, C. R., et al. (2017). Causes and consequences of movement: The interaction between foraging and landscape patterns (Doctoral dissertation).

Agostinelli, C., & Lund, U. (2023). R package circular: Circular statistics (version 0.5-0). https://CRAN.R-project.org/package=circular

Albert, A., Auffret, A. G., Cosyns, E., Cousins, S. A. O., D’hondt, B., Eichberg, C., Eycott, A. E., Heinken, T., Hoffmann, M., Jaroszewicz, B., Malo, J. E., Mårell, A., Mouissie, M., Pakeman, R. J., Picard, M., Plue, J., Poschlod, P., Provoost, S., Schulze, K. A., & Baltzinger, C. (2015). Seed dispersal by ungulates as an ecological filter: A trait-based meta-analysis. Oikos, 124(9), 1109–1120. 10.1111/oik.02512

Ashton, B. J., Kennedy, P., & Radford, A. N. (2020). Interactions with conspecific outsiders as drivers of cognitive evolution. Nature Communications, 11(1), 4937.

Avgar, T., Deardon, R., & Fryxell, J. M. (2013). An empirically parameterized individual based model of animal movement, perception, and memory. Ecological Modelling, 251, 158–172. 10.1016/j.ecolmodel.2012.12.002

Bagchi, R., Henrys, P. A., Brown, P. E., Burslem, D. F., Diggle, P. J., Gunatilleke, C. V., Gunatilleke, I. A., Kassim, A. R., Law, R., Noor, S., & Valencia, R. L. (2011). Spatial patterns reveal negative density dependence and habitat associations in tropical trees. Ecology, 92(9), 1723–1729. 10.1890/11-0335.1

Ban, S. D., Boesch, C., & Janmaat, K. R. (2014). Taï chimpanzees anticipate revisiting high-valued fruit trees from further distances. Animal Cognition, 17(6), 1353–1364. 10.1007/s10071-014-0771-y

Ban, S. D., Boesch, C., N’Guessan, A., N’Goran, E. K., Tako, A., & Janmaat, K. R. (2016). Taï chimpanzees change their travel direction for rare feeding trees providing fatty fruits. Animal Behaviour, 118, 135–147. 10.1016/j.anbehav.2016.05.014

Bartumeus, F., Campos, D., Ryu, W. S., Lloret-Cabot, R., Méndez, V., & Catalan, J. (2016). Foraging success under uncertainty: search tradeoffs and optimal space use. Ecology Letters, 19(11), 1299–1313. 10.1111/ele.12660

Batschelet, E. (1981). Circular Statistics in Biology. Academic Press.

Beisner, B. E., Haydon, D. T., & Cuddington, K. (2003). Alternative stable states in ecology. Frontiers in Ecology and the Environment, 1(7), 376–382.

Benhamou, S. (2014). Of scales and stationarity in animal movements. Ecology Letters, 17(3), 261–272. 10.1111/ele.12225

Benhamou, S., & Bovet, P. (1992). Distinguishing between elementary orientation mechanisms by means of path analysis. Animal Behaviour, 43(3), 371–377.

Benhamou, S., & Collet, J. (2015). Ultimate failure of the lévy foraging hypothesis: Two-scale searching strategies outperform scale-free ones even when prey are scarce and cryptic. Journal of theoretical biology, 387, 221–227.

Benhamou, S., & Poucet, B. (1995). A comparative analysis of spatial memory processes. Behavioural Processes, 35(1-3), 113–126.

Berger-Tal, O., & Bar-David, S. (2015). Recursive movement patterns: Review and synthesis across species. Ecosphere, 6(9), 1–12.

Bialozyt, R., Flinkerbusch, S., Niggemann, M., & Heymann, E. W. (2014). Predicting the seed shadows of a neotropical tree species dispersed by primates using an agent-based model with internal decision making for movements. Ecological Modelling, 278, 74–84.

Blake, S., Deem, S. L., Mossimbo, E., Maisels, F., & Walsh, P. (2009). Forest elephants: Tree planters of the congop. Biotropica, 41(4), 459–468. 10.1111/j.1744-7429.2009.00512.x

Blake, S., & Inkamba-Nkulu, C. (2004). Fruit, minerals, and forest elephant trails: Do all roads lead to Rome? Biotropica, 36(3), 392–401. 10.1111/j.1744-7429.2004.tb00332.x

Borah, B., & Beckman, N. G. (2022). Studying seed dispersal through the lens of movement ecology. Oikos, 2022(2).

Boyer, D., & Walsh, P. D. (2010). Modelling the mobility of living organisms in heterogeneous landscapes: Does memory improve foraging success? *Philosophical Transactions of the Royal Society A: Mathematical*, Physical and Engineering Sciences, 368(1933), 5645–5659. 10.1098/rsta.2010.0275

Bracis, C., Gurarie, E., Van Moorter, B., & Goodwin, R. A. (2015). Memory effects on movement behavior in animal foraging. PLoS ONE, 10(8), e0136057. 10.1371/journal.pone.0136057

Brehm, A. M., & Mortelliti, A. (2022). Small mammal personalities generate context dependence in the seed dispersal mutualism. Proceedings of the National Academy of Sciences, 119(15), e2113870119. 10.1073/pnas.2113870119

Burns, J. G., Foucaud, J., & Mery, F. (2011). Costs of memory: Lessons from ‘mini’brains. Proceedings of the Royal Society B: Biological Sciences, 278(1707), 923–929.

Burt, W. H. (1943). Territoriality and home range concepts as applied to mammals. Journal of mammalogy, 24(3), 346–352.

Calenge, C. (2006). The package adehabitat for the r software: Tool for the analysis of space and habitat use by animals. Ecological Modelling, 197, 1035.

Calenge, C., contributions from Stephane Dray, & Royer, M. (2023). Adehabitatlt: Analysis of animal movements [R package version 0.3.27]. https://CRAN.R-project.org/package=adehabitatLT

Callaway, R. M. (1995). Positive interactions among plants. The Botanical Review, 61, 306–349.

Campos-Arceiz, A., & Blake, S. (2011). Megagardeners of the forest - the role of elephants in seed dispersal. Acta Oecologica, 37(6), 542–553. 10.1016/j.actao.2011.01.014

Chapman, C. A., Bonnell, T. R., Gogarten, J. F., Lambert, J. E., Omeja, P. A., Twinomugisha, D., Wasserman, M. D., & Rothman, J. M. (2013). Are primates ecosystem engineers? International Journal of Primatology, 34(1), 1–14. 10.1007/s10764-012-9645-9

Chuine, I., Belmonte, J., & Mignot, A. (2000). A modelling analysis of the genetic variation of phenology between tree populations. Journal of Ecology, 88(4), 561–570. Retrieved June 21, 2023, from http://www.jstor.org/stable/2648589

Clark, J. S., Silman, M., Kern, R., Macklin, E., & HilleRisLambers, J. (1999). Seed dispersal near and far: Patterns across temperate and tropical forests. Ecology, 80(5), 1475–1494.

Connell, J. H. (1971). On the role of natural enemies in preventing competitive exclusion in some marine animals and in rain forest trees. Dynamics of populations, 298(312).

Côrtes, M. C., & Uriarte, M. (2013). Integrating frugivory and animal movement: A review of the evidence and implications for scaling seed dispersal. Biological Reviews, 88(2), 255–272.

Craine, J. M., & Dybzinski, R. (2013). Mechanisms of plant competition for nutrients, water and light. Functional Ecology, 27(4), 833–840. 10.1111/1365-2435.12081

de Guinea, M., Estrada, A., Nekaris, K. A.-I., & Van Belle, S. (2019). Arboreal route navigation in a Neotropical mammal: Energetic implications associated with tree monitoring and landscape attributes. Movement Ecology, 7(1), 1–12. 10.1186/s40462-019-0187-z

Di Fiore, A., & Suarez, S. A. (2007). Route-based travel and shared routes in sympatric spider and woolly monkeys: Cognitive and evolutionary implications. Animal cognition, 10, 317–329.

Doran-Sheehy, D. M., Greer, D., Mongo, P., & Schwindt, D. (2004). Impact of ecological and social factors on ranging in western gorillas. American Journal of Primatology: Official Journal of the American Society of Primatologists, 64(2), 207–222.

Eddelbuettel, D. (2013). Seamless R and C++ integration with Rcpp [ISBN 978-1-4614-6867-7]. Springer. 10.1007/978-1-4614-6868-4

Eddelbuettel, D., Francois, R., Allaire, J., Ushey, K., Kou, Q., Russell, N., Ucar, I., Bates, D., & Chambers, J. (2023). Rcpp: Seamless r and c++ integration [R package version 1.0.11]. https://CRAN.R-project.org/package=Rcpp

Eddelbuettel, D., & François, R. (2011). Rcpp: Seamless R and C++ integration. Journal of Statistical Software, 40(8), 1–18. 10.18637/jss.v040.i08

Fagan, W. F., Lewis, M. A., Auger-Méthé, M., Avgar, T., Benhamou, S., Breed, G., Ladage, L., Schlägel, U. E., Tang, W. W., Papastamatiou, Y. P., Forester, J., & Mueller, T. (2013). Spatial memory and animal movement. Ecology Letters, 16(10), 1316–1329. 10.1111/ele.12165

Flörchinger, M., Braun, J., Böhning-Gaese, K., & Schaefer, H. M. (2010). Fruit size, crop mass, and plant height explain differential fruit choice of primates and birds. Oecologia, 164(1), 151–161. 10.1007/s00442-010-1655-8

Fragoso, J. M. V. (1997). Tapir-generated seed shadows: Scale-dependent patchiness in the amazon rain forest. Journal of Ecology, 85(4), 519–529.

Fragoso, J. M. V., Silvius, K. M., & Correa, J. A. (2003). Long-distance seed dispersal by tapirs increases seed survival and aggregates tropical trees. Ecology, 84(8), 1998–2006. 10.1890/01-0621

Fryxell, J. M., Hazell, M., Börger, L., Dalziel, B. D., Haydon, D. T., Morales, J. M., McIntosh, T., & Rosatte, R. C. (2008). Multiple movement modes by large herbivores at multiple spatiotemporal scales. Proceedings of the National academy of Sciences, 105(49), 19114–19119.

Fuzessy, L., Balbuena, J. A., Nevo, O., Tonos, J., Papinot, B., Park, D., Sol, D., Valenta, K., Razafindratsima, O., & Verdu, M. (2023). Friends or foes? plant-animal coevolutionary history is driven by both mutualistic and antagonistic interactions. BioRxiv.

Fuzessy, L., Silveira, F. A., Culot, L., Jordano, P., & Verdú, M. (2022a). Phylogenetic congruence between Neotropical primates and plants is driven by frugivory. Ecology Letters, 25(2), 320–329. 10.1111/ele.13918

Fuzessy, L., Sobral, G., & Culot, L. (2022b). Linking howler monkey ranging and defecation patterns to primary and secondary seed dispersal. American Journal of Primatology, 84(2), e23354. 10.1002/ajp.23354

Fuzessy, L. F., Janson, C., & Silveira, F. A. (2018). Effects of seed size and frugivory degree on dispersal by neotropical frugivores. Acta Oecologica, 93, 41–47.

Gelmi-Candusso, T. A., Heymann, E. W., & Heer, K. (2017). Effects of zoochory on the spatial genetic structure of plant populations. Molecular Ecology, 26(21), 5896–5910. 10.1111/mec.14351

Gómez, J. M., & Verdú, M. (2012). Mutualism with plants drives primate diversification. Systematic Biology, 61(4), 567–577. 10.1093/sysbio/syr127

Gottsberger, G., & Silberbauer-Gottsberger, I. L. S. E. (1983). Dispersal and distribution in the cerrado vegetation of Brazil. Sonderbd natuwiss. ver. Hamburg, 7, 315–352.

Green, A. J., Baltzinger, C., & Lovas-Kiss, Á. (2022). Plant dispersal syndromes are unreliable, especially for predicting zoochory and long-distance dispersal. Oikos, 2022(2).

Grove, M. (2013). The evolution of spatial memory. Mathematical Biosciences, 242(1), 25–32. 10.1016/j.mbs.2012.11.011

Hansen, M. J., Schaerf, T. M., & Ward, A. J. (2015). The effect of hunger on the exploratory behaviour of shoals of mosquitofish gambusia holbrooki. Behaviour, 152(12-13), 1659–1677.

Havmøller, L. W., Loftus, J. C., Havmøller, R. W., Alavi, S. E., Caillaud, D., Grote, M. N., Hirsch, B. T., Tórrez-Herrera, L. L., Kays, R., & Crofoot, M. C. (2021). Arboreal monkeys facilitate foraging of terrestrial frugivores. Biotropica, 53(6), 1685–1697.

Hoyle, A., Bowers, R. G., White, A., & Boots, M. (2008). The influence of trade-off shape on evolutionary behaviour in classical ecological scenarios. Journal of Theoretical Biology, 250(3), 498– 511.

Hubbell, S. P. (1979). Tree dispersion, abundance, and diversity in a tropical dry forest: That tropical trees are clumped, not spaced, alters conceptions of the organization and dynamics. Science, 203(4387), 1299–1309.

Irwin, M. T., Samonds, K. E., Raharison, J.-L., & Wright, P. C. (2004). Lemur Latrines: Observations of Latrine Behavior in Wild Primates and Possible Ecological Significance. Journal of Mammalogy, 85(3), 420–427. 10.1644/1383937

Janmaat, K. R., Ban, S. D., & Boesch, C. (2013). Taï chimpanzees use botanical skills to discover fruit: What we can learn from their mistakes. Animal Cognition, 16(6), 851–860. 10.1007/s10071-013-0617-z

Janmaat, K. R., Byrne, R. W., & Zuberbühler, K. (2006). Primates take weather into account when searching for fruits. Current Biology, 16(12), 1232–1237. 10.1016/j.cub.2006.04.031

Janmaat, K. R., Chapman, C. A., Meijer, R., & Zuberbühler, K. (2012). The use of fruiting synchrony by foraging mangabey monkeys: A ‘simple tool’ to find fruit. Animal Cognition, 15(1), 83–96. 10.1007/s10071-011-0435-0

Janson, C. H. (1998). Experimental evidence for spatial memory in foraging wild capuchin monkeys, Cebus apella. Animal Behaviour, 55(5), 1229–1243. 10.1006/anbe.1997.0688

Janson, C. H. (2016). Capuchins, space, time and memory: An experimental test of what-where-when memory in wild monkeys. Proceedings of the Royal Society B: Biological Sciences, 283(1840), 20161432. 10.1098/rspb.2016.1432

Janzen, D. H. (1970). Herbivores and the number of tree species in tropical forests. The American Naturalist, 104(940), 501–528.

John, E. A., Francesca, S., Burman, O. H. P., Anna, W., & Pike, T. W. (2016). Plant ecology meets animal cognition: impacts of animal memory on seed dispersal. Plant Ecology, 217, 1441–1456. 10.1007/s11258-016-0652-3

Jordano, P., García, C., Godoy, J. A., & García-Castaño, J. L. (2007). Differential contribution of frugivores to complex seed dispersal patterns. Proceedings of the National Academy of Sciences, 104(9), 3278–3282. 10.1073/pnas.0606793104

Kraemer, P. J., & Golding, J. M. (1997). Adaptive forgetting in animals. Psychonomic Bulletin & Review, 4, 480–491.

Lambert, J. E., & Garber, P. A. (1998). Evolutionary and ecological implications of primate seed dispersal. American Journal of primatology, 45(1), 9–28.

Lamontagne, J. M., & Boutin, S. (2007). Local-scale synchrony and variability in mast seed production patterns of picea glauca. Journal of Ecology, 95(5), 991–1000.

LaScala-Gruenewald, D. E., Mehta, R. S., Liu, Y., & Denny, M. W. (2019). Sensory perception plays a larger role in foraging efficiency than heavy-tailed movement strategies. Ecological Modelling, 404, 69–82. 10.1016/j.ecolmodel.2019.02.015

Lloyd, M. (1967). Mean crowding. The Journal of Animal Ecology, 1–30. 10.2307/3012

Lomáscolo, S. B., & Schaefer, H. M. (2010). Signal convergence in fruits: A result of selection by frugivores? Journal of Evolutionary Biology, 23(3), 614–624. 10.1111/j.1420-9101.2010.01931.x

Lyngdoh, N., Gunaga, R., Joshi, G., Vasudeva, R., Ravikanth, G., & Shaanker, R. U. (2012). Influence of geographic distance and genetic dissimilarity among clones on flowering synchrony in a teak (*Tectona grandis Linn. f*) clonal seed orchard. Silvae Genetica, 61(1-6), 10–18.

Masi, S., Cipolletta, C., & Robbins, M. M. (2009). Western lowland gorillas (*Gorilla gorilla gorilla*) change their activity patterns in response to frugivory. American Journal of Primatology: Official Journal of the American Society of Primatologists, 71(2), 91–100.

Milton, K. (1981). Distribution Patterns of Tropical Plant Foods as an Evolutionary Stimulus to Primate Mental Development. American Anthropologist, 83(3), 534–548. 10.1525/aa.1981.83.3.02a00020

Morales, J. M., & Moran Lopez, T. (2022). Mechanistic models of seed dispersal by animals. Oikos, 2022(2).

Morales, J. M., & Morán López, T. (2022). Mechanistic models of seed dispersal by animals. Oikos, 2022(2). 10.1111/oik.08328

Nathan, R., Getz, W. M., Revilla, E., Holyoak, M., Kadmon, R., Saltz, D., & Smouse, P. E. (2008). A movement ecology paradigm for unifying organismal movement research. Proceedings of the National Academy of Sciences, 105(49), 19052–19059.

Paradis, E., & Schliep, K. (2019). Ape 5.0: An environment for modern phylogenetics and evolutionary analyses in R. Bioinformatics, 35, 526–528. 10.1093/bioinformatics/bty633

Pearce, F., Carbone, C., Cowlishaw, G., & Isaac, N. J. (2013). Space-use scaling and home range overlap in primates. Proceedings of the Royal Society B: Biological Sciences, 280(1751), 20122122.

Post, D. M., & Palkovacs, E. P. (2009). Eco-evolutionary feedbacks in community and ecosystem ecology: Interactions between the ecological theatre and the evolutionary play. Philosophical Transactions of the Royal Society B: Biological Sciences, 364(1523), 1629–1640.

Pouvelle, S., Jouard, S., Feer, F., Tully, T., & Ponge, J.-F. (2009). The latrine effect: Impact of howler monkeys on the distribution of small seeds in a tropical rain-forest soil. Journal of Tropical Ecology, 25(3), 239–248. 10.1017/S0266467409005987

Presotto, A., Verderane, M. P., Biondi, L., Mendonça-Furtado, O., Spagnoletti, N., Madden, M., & Izar, P. (2018). Intersection as key locations for bearded capuchin monkeys (*Sapajus libidinosus*) traveling within a route network. Animal Cognition, 21(3), 393–405. 10.1007/s10071-018-1176-0

Pringle, R. M. (2008). Elephants as agents of habitat creation for small vertebrates at the patch scale. Ecology, 89(1), 26–33. 10.1890/07-0776.1

R Core Team. (2022). R: A language and environment for statistical computing. R Foundation for Statistical Computing. Vienna, Austria. https://www.R-project.org/

Raichle, M. E. (2006). The brain’s dark energy. Science-New York Then Washington*-*, 314(5803), 1249.

Remis, M. J. (1997). Western lowland gorillas (*Gorilla gorilla gorilla*) as seasonal frugivores: Use of variable resources. American Journal of Primatology, 43(2), 87–109.

Riotte-Lambert, L., Benhamou, S., & Chamaillé-Jammes, S. (2015). How memory-based movement leads to nonterritorial spatial segregation. American Naturalist, 185(4), E103–E116. 10.1086/680009

Riotte-Lambert, L., Benhamou, S., & Chamaillé-Jammes, S. (2017). From randomness to traplining: a framework for the study of routine movement behavior. Behavioral Ecology, 28(1), 280–287. 10.1093/beheco/arw154

Riotte-Lambert, L., & Matthiopoulos, J. (2020). Environmental predictability as a cause and consequence of animal movement. Trends in Ecology and Evolution, 35(2), 163–174. 10.1016/j.tree.2019.09.009

Robira, B., Benhamou, S., Bayanga, E. O., Breuer, T., & Masi, S. (2023a). How do primates decide where to feed? insights from wild western gorillas. Animal behaviour.

Robira, B., Benhamou, S., Fuh, T. N., & Masi, S. (2023b). Do seasonal frugivory and cognition shape foraging movements in wild western gorillas? In Movement ecology of afrotropical forest mammals (pp. 151–170). Springer.

Robira, B., Benhamou, S., Masi, S., Llaurens, V., & Riotte-Lambert, L. (2021). Foraging efficiency in temporally predictable environments: Is a long-term temporal memory really advantageous? Royal Society Open Science, 8(9), 210809. 10.1098/rsos.210809

Robira, B., & Perez-Lamarque, B. (2023). Primate sympatry shapes the evolution of their brain architecture. Peer Community Journal, 3.

Rogers, H. S., Beckman, N. G., Hartig, F., Johnson, J. S., Pufal, G., Shea, K., Zurell, D., Bullock, J. M., Cantrell, R. S., Loiselle, B., et al. (2019). The total dispersal kernel: A review and future directions. AoB Plants, 11(5), plz042.

Rojas, T. N., Bruzzone, O. A., Zampini, I. C., Isla, M. I., & Blendinger, P. G. (2021). A combination of rules govern fruit trait preference by frugivorous bat and bird species: Nutrients, defence and size. Animal Behaviour, 176, 111–123.

Rosati, A. G. (2017). Foraging cognition: Reviving the Ecological Intelligence Hypothesis. Trends in Cognitive Sciences, 21(9), 691–702. 10.1016/j.tics.2017.05.011

Russo, S. E., Portnoy, S., & Augspurger, C. K. (2006a). Incorporating animal behavior into seed dispersal models: Implications for seed shadows. Ecology, 87(12), 3160–3174.

Russo, S. E., Portnoy, S., & Augspurger, C. K. (2006b). Incorporating animal behaviour into seed dispersal models: Implications for seed shadows. Ecology, 87(12), 3160–3174. 10.1890/0012-9658(2006)87[3160:IABISD]2.0.CO;2

Sakamoto, Y., & Takatsuki, S. (2015). Seeds Recovered from the Droppings at Latrines of the Raccoon Dog (*Nyctereutes procyonoides viverrinus*): The Possibility of Seed Dispersal. Zoological Science, 32(2), 157–162. 10.2108/zs140107

Sassi, Y., Nouzieres, B., Scacco, M., Tremblay, Y., Duriez, O., & Robira, B. (in press). The use of social information in vulture flight decisions. Proceedings of the Royal Society B: Biological Sciences.

Schmal, C., Myung, J., Herzel, H., & Bordyugov, G. (2017). Moran’s i quantifies spatio-temporal pattern formation in neural imaging data. Bioinformatics, 33(19), 3072–3079.

Seidler, T. G., & Plotkin, J. B. (2006). Seed dispersal and spatial pattern in tropical trees. PLoS Biology, 4(11), 2132–2137. 10.1371/journal.pbio.0040344

Shanahan, M., So, S., Compton, S. G., & Corlett, R. (2001). Fig-eating by vertebrate frugivores: A global review. Biological reviews, 76(4), 529–572.

Soldati, F., et al. (2015). Animal cognition meets ecosystem ecology: The impact of cognition on seed dispersal (Doctoral dissertation). University of Lincoln.

Sussman, R. W. (1991). Primate origins and the evolution of angiosperms. American Journal of Primatology, 23(4), 209–223. 10.1002/ajp.1350230402

Teichroeb, J. A., & Vining, A. Q. (2019). Navigation strategies in three nocturnal lemur species: diet predicts heuristic use and degree of exploratory behavior. Animal Cognition, 22(3), 343–354. 10.1007/s10071-019-01247-4

Tello-Ramos, M. C., Branch, C. L., Kozlovsky, D. Y., Pitera, A. M., & Pravosudov, V. V. (2019). Spatial memory and cognitive flexibility trade-offs: To be or not to be flexible, that is the question. Animal Behaviour, 147, 129–136.

Terborgh, J. (1986). Community aspects of frugivory in tropical forests. In Frugivores and seed dispersal (pp. 371–384). Springer.

Tiffney, B. H., & Mazer, S. J. (1995). Angiosperm growth habit, dispersal and diversification reconsidered. Evolutionary Ecology, 9, 93–117.

Trapanese, C., Meunier, H., & Masi, S. (2019a). What, where and when: spatial foraging decisions in primates. Biological Reviews, 94(2), 483–502. 10.1111/brv.12462

Trapanese, C., Meunier, H., & Masi, S. (2022). Do primates flexibly use spatio-temporal cues when foraging? Quarterly Journal of Experimental Psychology, 75(2), 232–244. 10.1177/1747021820970724

Trapanese, C., Robira, B., Tonachella, G., Di Gristina, S., Meunier, H., & Masi, S. (2019b). Where and what? Frugivory is associated with more efficient foraging in three semi-free ranging primate species. Royal Society Open Science, 6(5), 181722. 10.1098/rsos.181722

Van Woerden, J. T., Van Schaik, C. P., & Isler, K. (2010). Effects of seasonality on brain size evolution: Evidence from strepsirrhine primates. The American Naturalist, 176(6), 758–767.

Vanleeuwe, H., & Gautier-Hion, A. (1998). Forest elephant paths and movements at the Odzala National Park, Congo: The role of clearings and Marantaceae forests. African Journal of Ecology, 36(2), 174–182. 10.1046/j.1365-2028.1998.00123.x

van Woerden, J. T., Willems, E. P., van Schaik, C. P., & Isler, K. (2012). Large brains buffer energetic effects of seasonal habitats in catarrhine primates. Evolution, 66(1), 191–199.

Vittoz, P., & Engler, R. (2007). Seed dispersal distances: A typology based on dispersal modes and plant traits. Botanica Helvetica, 117(2), 109–124. 10.1007/s00035-007-0797-8

Vuarin, P., & Henry, P.-Y. (2014). Field evidence for a proximate role of food shortage in the regulation of hibernation and daily torpor: A review. Journal of Comparative Physiology B, 184, 683–697.

Warren, W. H. (2019). Non-euclidean navigation. Journal of Experimental Biology, 222.

Westcott, D. A., Bentrupperbäumer, J., Bradford, M. G., & McKeown, A. (2005). Incorporating patterns of disperser behaviour into models of seed dispersal and its effects on estimated dispersal curves. Oecologia, 146, 57–67.

White, J. W., Rassweiler, A., Samhouri, J. F., Stier, A. C., & White, C. (2014). Ecologists should not use statistical significance tests to interpret simulation model results. Oikos, 123(4), 385–388.

Zuberbühler, K., & Janmaat, K. (2010). Foraging cognition in non-human primates. Primate neuroethology, 64–83.

Zwolak, R. (2018). How intraspecific variation in seed-dispersing animals matters for plants. Biological Reviews, 93(2), 897–913. 10.1111/brv.12377

Zwolak, R., & Sih, A. (2020). Animal personalities and seed dispersal: A conceptual review. Functional Ecology, 34(7), 1294–1310. 10.1111/1365-2435.13583

